# Induction of cortical ON/OFF periods in awake mice fulfills sleep functions

**DOI:** 10.1101/2025.10.04.680459

**Authors:** Kort Driessen, Fabio Squarcio, Giulio Tononi, Chiara Cirelli

## Abstract

Can animals obtain core benefits of sleep while remaining awake? In mammals, slow-wave sleep is characterized by synchronized neuronal activity alternating between ON and OFF periods. Slow-wave activity and synchrony reflect sleep need, are correlated with synaptic strength in cortical circuits, and promote synaptic down-selection and memory consolidation. To address the above question, we locally induced alternating ON/OFF periods during wakefulness using optogenetics in mice. This led to a local, ipsilateral reduction of slow-wave activity and synchrony during subsequent sleep and to reduced markers of synaptic strength. Moreover, bilateral induction of OFF periods over sensorimotor cortex during sleep deprivation restored memory consolidation. Thus, inducing ON/OFF activity during wakefulness is sufficient to reduce local sleep need and fulfills core functions of sleep.

## Main Text

Sleep is a universal behavioral state typically characterized by quiescence and decreased responsiveness to stimuli. It is also homeostatically regulated, becoming deeper and often longer after extended periods of wakefulness^1^. Sleep is thought to perform essential functions that the brain cannot carry out when connected to the environment^2–4^. Chief among these is synaptic homeostasis, which prevents synaptic saturation, restores the ability to learn^5–7^, and promotes memory consolidation^5–8^.

Non-rapid eye movement (NREM) sleep, which accounts for ∼80% of sleep in adults, has been closely associated with synaptic homeostasis^9,10^ and associated memory functions^11–13^. A defining feature of NREM sleep is the synchronized, low frequency alternation between periods of high population spiking activity of cortical neurons (ON periods) and periods of population silence (OFF periods)^14–16^. This widespread, synchronized, bistable activity pattern underlies the occurrence of slow waves in recordings of cortical field potentials such as the electroencephalogram (EEG). Slow-wave activity (SWA), the EEG power in the delta range during NREM sleep (0.5-4 Hz), is a well-established marker of sleep need and homeostasis, increasing after sleep deprivation and learning and declining during sleep in both rodents and humans^1^. There is substantial evidence, in both animals and humans, that the synchronous ON/OFF activity patterns that underlie slow waves in NREM sleep may be key mechanistic mediators of sleep’s restorative functions^9,10,13,17–21^. This raises the question whether, if one could induce synchronous ON/OFF activity patterns locally in individuals who are otherwise awake, some of the homeostatic functions of sleep and associated benefits may be carried out without disconnecting from the environment. Indeed, an evolutionary proof of concept is offered by some cetaceans and bird species that can enter NREM sleep with one hemisphere while the other hemisphere remains vigilant^22^.

To address this question experimentally, we developed a method to locally induce NREM sleep-like ON/OFF patterns in one hemisphere of awake mice using complementary optogenetic approaches, while recording unit activity from homotopic networks in both hemispheres. As shown below, this resulted in reduced SWA and synchrony in subsequent NREM sleep on the stimulated side. We further show that a broad induction of NREM sleep-like cortical ON/OFF periods in awake animals renormalizes molecular markers of excitatory synaptic strength and improves memory consolidation, thus fulfilling key homeostatic functions of sleep in awake, behaving animals.

### Recording and manipulation of homotopic cortical networks

We implanted adult mice with bilateral linear silicon probes, which allowed us to record laminar multi-unit activity (MUA) and local field potential (LFP) in local cortical networks of homotopic brain regions. An optic ferrule was attached to one of the probes (hereafter termed the ‘optrode’) so that the cortical network recorded by that probe could be optogenetically manipulated independently of its homotopic partner (hereafter termed the ‘contralateral control’ probe, Fig. 1a). We recorded LFP and MUA data from the bilateral silicon probe implant, as well as electromyogram (EMG), continuously for 48 hours, and determined whether the mouse was awake, in NREM sleep, or in REM sleep for the duration of the recording (Fig. 1b,c). Beginning at light onset, mice were sleep deprived for 5 hours by introducing novel objects into the home cage. This sleep deprivation (SD) protocol reliably kept mice awake for the full 5-hour period (Extended Data Fig. 1) and reliably resulted in a large “rebound” in SWA in subsequent NREM sleep which then declined as the mouse slept – a clear demonstration of SWA reflecting the buildup and subsequent dissipation of the pressure to sleep (Fig. 1b). In control conditions, measures of sleep pressure such as SWA were tightly coupled in both probes (Fig. 1b,c).

**Fig. 1.**
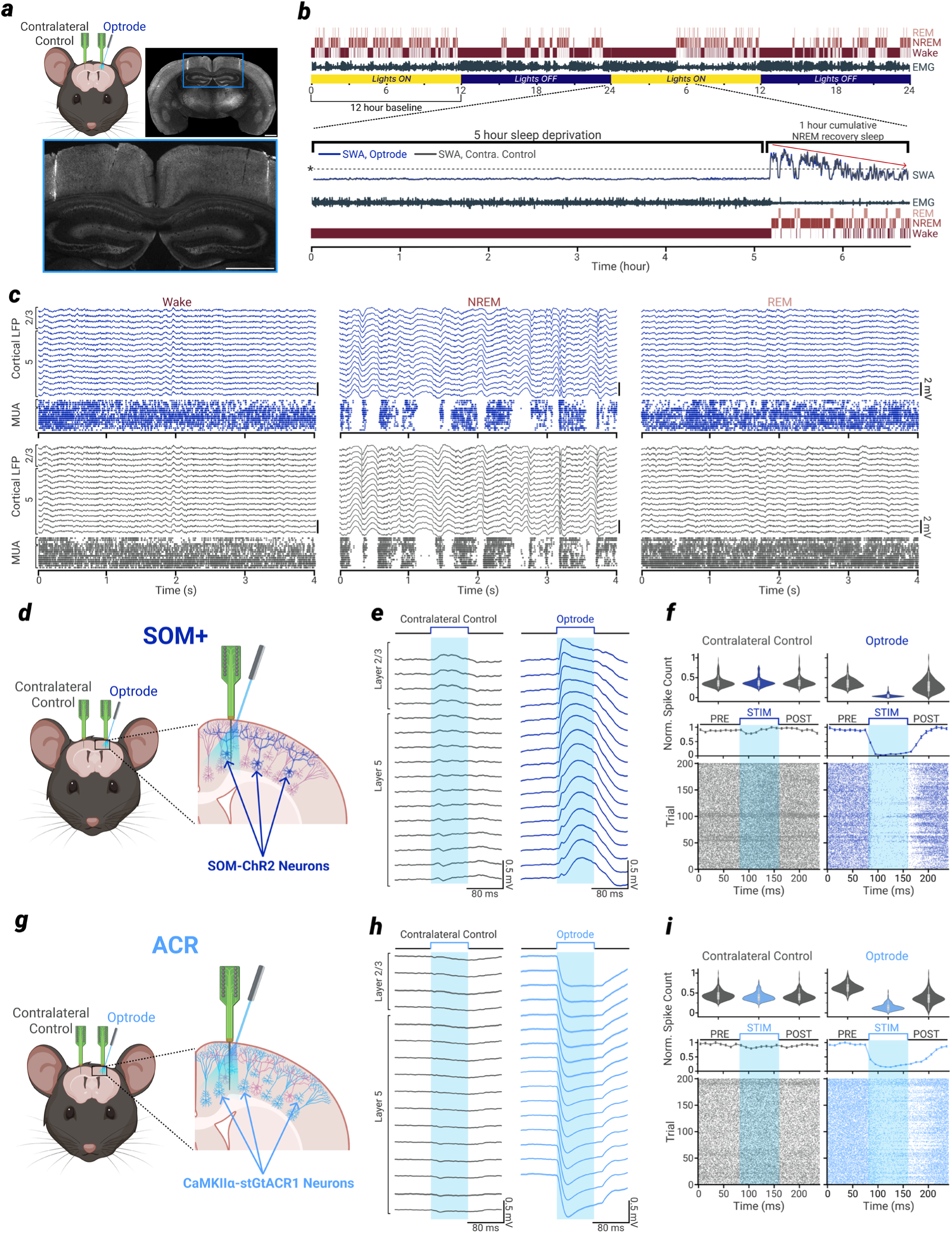
Chronic recording and optogenetic manipulation of local cortical networks **a**, Top left: implant schematic. Two silicon probes are inserted to the same coordinates exactly contralateral (i.e. homotopic) to each other. One probe is connected to an optic ferrule for the delivery of light in optogenetic experiments (‘Optrode’) and the other has no attached ferrule and can only record (‘Contralateral Control’). Top right: representative coronal section showing the fluorescent tracks where the probes were implanted. Area in blue box is enlarged at bottom. Scale bars represent 1 mm. **b**, Upper: example hypnogram and corresponding muscular activity (electromyogram, EMG) from a continuous 48-hour recording. Lower: sleep deprivation (SD) via novel objects for 5 hours, beginning at lights-on. Recovery sleep is defined as the first 1 hour of cumulative NREM sleep after SD. Slow wave activity (SWA, spectral power between 0.5 - 4 Hz) for optrode and contralateral control probe is expressed relative to the NREM average during the 12-hour baseline. Note the tight SWA correspondence between the two probes. *Dotted line indicates the 12-hour NREM baseline SWA mean. Red arrow indicates the homeostatic decline in SWA as the mouse spends time asleep. **c**, Example local field potential (LFP) and multi-unit activity (MUA) data in Wake, NREM sleep, REM sleep from a mouse with probes implanted in secondary motor area (M2). Layer boundaries across the 16 channels are shown for both probes at far left. **d**, SOM+ mice express ChR2 in somatostatin-producing interneurons. The optogenetic activation of these neurons (dark blue cells indicated by arrows) leads to inhibition of all neuron types in the cortical network. **e**, Example SOM+ mouse average field potential response to 200 80ms light pulses. Data represent mean ± SEM. Data is from one representative mouse with implant in M2. **f**, Bottom: raster plot with each row showing MUA collapsed across all channels for one 80ms light pulse. Middle: spike count in 10ms bins normalized to maximum value, data represent mean ± SEM. Top: spike counts during each full 80ms epoch (pre-stim, stim, post-stim) normalized to maximum value. **g**, ACR mice express soma-targeted Guillardia theta anion-conducting channelrhodopsin-1 (stGtACR1) in CaMKIIα-producing neurons (mainly pyramidal neurons, light blue cells indicated by arrows). When stGtACR1 is optogenetically activated, all CaMKIIα neurons in the cortical network are inhibited. **h**, Same as in **e**, but for one representative ACR mouse with an implant located in M2. **i**, Same as in **f**, but for the ACR mouse shown in **h**.

We used two optogenetic mouse models to induce NREM-sleep like patterns of ON/OFF activity in one local cortical network independently of neurons in the homotopic cortical network Extended Data Movie 1). One strategy involved recruiting somatostatin-expressing neurons, a class of cortical interneurons known to play a key role in the generation of sleep slow waves^23^. A complementary approach involved directly inhibiting excitatory pyramidal neurons.

In the first model, we crossed mice carrying a transgene encoding channelrhodopsin-2 (ChR2) with mice expressing the enzyme Cre under the control of the somatostatin promoter (Fig. 1d). In the generated mouse line (hereafter termed ‘SOM+’ mice) somatostatin-expressing neurons could be optogenetically activated, resulting in positive-going, NREM sleep-like slow waves (Fig. 1e) in the LFP, and the associated inhibition of spiking activity in nearly all other neurons in the local cortical network recorded by the optrode (Fig. 1f).

In the second model, we crossed mice expressing soma-targeted *Guillardia theta* anion-conducting channelrhodopsin-1 (stGtACR1^24^) with mice expressing the enzyme Cre under the control of the CaMKIIα promoter (Fig. 1g). In the generated mouse line (hereafter termed ‘ACR’ mice) optogenetic stimulation led to potent inhibition of nearly all cortical neurons, resulting in negative-going slow waves (Fig. 1h), and the associated inhibition of nearly all spiking activity in the local cortical network recorded by the optrode (Fig. 1i).

### Induction of OFF periods during wakefulness reduces SWA in subsequent NREM sleep

We first wondered whether the sustained induction of NREM-like OFF periods in a local cortical network of an awake mouse could reduce SWA only in that network in subsequent NREM sleep. To test this hypothesis, we performed a 5-hour SD via novel objects as described in Fig. 1b but, starting 30 minutes before the end of the SD, we induced OFF periods similar to those seen in NREM sleep (see materials and methods), while the SD continued and the mouse remained awake (video S1). In both SOM+ mice (N = 10) and ACR mice (N = 9), the induction of OFF periods led to a large increase in SWA during the stimulation period on the optrode side, and not on the homotopic contralateral control side (Fig. 2a-c, Extended Data Fig.2a,b).

**Fig. 2.**
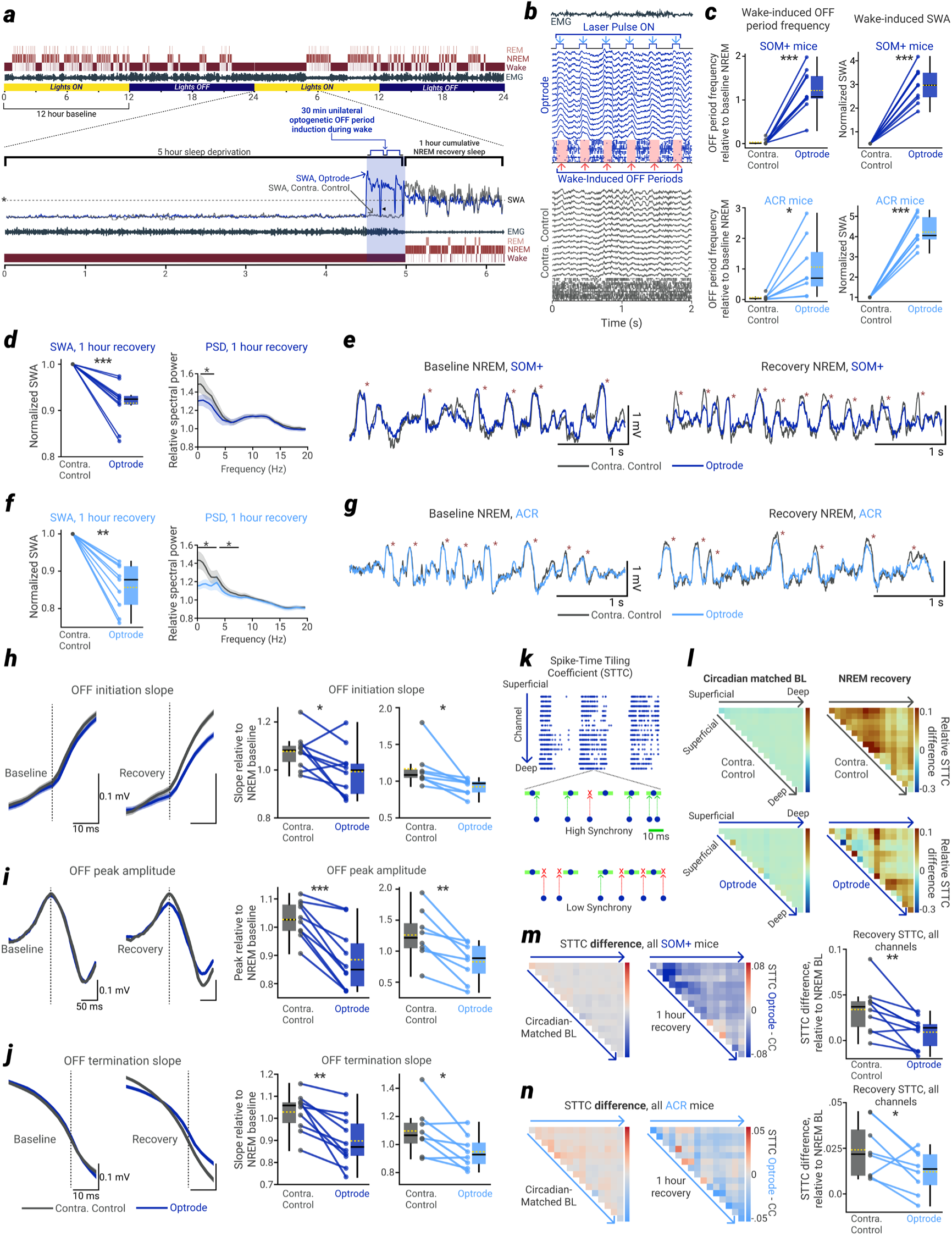
OFF period induction during wake reduces SWA and synchrony in subsequent sleep **a**, Experiment overview; in the last 30 minutes at the end of 5 hours of SD, OFF periods are optogenetically induced. SWA for optrode and contralateral control probe, EMG, and hypnogram are shown across the duration of the SD experiment and recovery sleep. Dashed line indicates the SWA mean over the 12 hour NREM baseline (indicated at left by an asterisk). Black triangle indicates the first 45 second break in between bouts of OFF induction (see methods). **b**, LFP traces and MUA during OFF period induction for an example optrode (top, blue) and contralateral control probe (bottom, gray) from one representative mouse. Note the tight correspondence between laser pulses (blue arrows at top) and algorithmically-detected OFF periods (red arrows, bottom). **c**, OFF period frequency (left) and SWA (right) during the induction period, relative to the 12-hour baseline NREM mean, for all SOM+ mice (top, dark blue) and ACR mice (bottom, light blue). SWA is normalized to the contralateral control probe. Dotted gold line represents mean in all boxplots. **d**, Left: Average SWA in 1 hour of cumulative NREM sleep, averaged across all channels of all SOM+ animals. Values are calculated relative to the 12-hour baseline NREM mean and then normalized to the contralateral control probe for visualization. Right: Average power spectral density (PSD) during 1 hour of cumulative NREM sleep, across all channels of all SOM+ animals. Values are relative to the 12-hour baseline NREM mean, shading represents ±SEM. **e**, Example NREM traces from a representative SOM+ animal in baseline and recovery sleep. Asterisks mark slow waves. Note the tight correspondence in baseline, and dissociation in recovery. **f**, Same as in **d**, but for all ACR animals **g**, Same as in **e**, but for a representative ACR animal **h**, Left: example traces from one representative SOM+ animal showing field potential slopes at OFF period initiation; data show an average ± SEM of traces from the first 1500 OFF periods in baseline and recovery sleep. Vertical dotted line marks OFF period start. Right: average field potential slope at OFF period initiation across all SOM+ and ACR mice. Data in **h-j** are for the first 30 minutes of the NREM recovery, when values were most reliably above baseline. **i**, Same as in **h**, but for the peak OFF period amplitude. Data are aligned to field potential peak during the OFF period, shown by vertical dotted line. **j**, Same as in **h**, but for the slope at OFF period termination. Vertical dotted line marks OFF period end. **k**, Spike-time tiling coefficient (STTC), a rate-independent measure of neural synchrony, is calculated for a pair of spike trains. The measure is high when a large proportion of spikes from train-1 have at least one other spike from train-2 within ±5 ms (light green boxes). In the ‘high synchrony’ example, 4/5 spikes from train-1 have at least one partner spike in train-2 within ±5 ms, whereas in the ‘low synchrony’ example, only 1/5 spikes have a partner. **l**, STTC is calculated across all unique channel-pairs for each probe of each mouse. Shown is the full STTC matrix for one representative SOM+ mouse’s contralateral control probe (top) and optrode (bottom) for the 1-hour NREM recovery period (right) and the circadian-matched NREM period from the baseline (left). Values are expressed as a difference between the condition mean and the 12-hour baseline NREM mean. **m**, Left: the difference matrix (optrode – contralateral control) averaged across all SOM+ mice for the 1-hour NREM recovery period (right) and the circadian-matched period from the baseline (left). Right: all-channel STTC average across all SOM+ mice for the 1-hour NREM recovery period relative to the full 12 hour baseline NREM mean. **n**, Same as in **m**, but for ACR mice. In all panels, *P<.05 | **P<.005 | ***P<.0005 | n.s. P>.05.

In the first hour of NREM sleep following SD, SWA recorded by the optrode was reduced relative to SWA on the contralateral control probe, in both SOM+ (Fig. 2d) and ACR (Fig. 2f) mice. This difference is also evident in the raw traces from matched channels of the same animal, where an extremely tight correspondence in slow wave magnitude and the general structure of the LFP is present during baseline NREM sleep but not during recovery NREM sleep (Fig. 2e,g). Furthermore, the reduction in SWA on the optrode recovered to equivalence with the contralateral control probe within a few hours of NREM sleep (Extended Data Fig.2c) and was specific for slow frequencies (i.e. was not a broadband reduction, Fig. 2d,f, right), not extending to other frequency bands tested (except the theta band for ACR mice, Extended Data Fig.2d).

When we performed the same OFF period induction protocol in wildtype control mice (N = 5), we did not see any change in overall firing rate, nor an increased frequency of OFF periods on the optrode relative to the contralateral control (Extended Data Fig. 3a). During the 1-hour NREM recovery period, we also observed no change in SWA, firing rate, or a general change in power spectral density (Extended Data Fig. 3b). Thus, the local induction of OFF periods during wakefulness leads to a local reduction of sleep pressure, as indexed by local SWA.

### Induction of OFF periods during wakefulness reduces network synchrony in subsequent NREM sleep

Next, we investigated the mechanisms underlying the decline of SWA in recovery sleep following OFF period induction during wake. We first determined that the frequency and duration of OFF periods during the first 1 hour of recovery sleep did not differ between the optrode side and the contralateral control (materials and methods, Extended Data Fig. 4a,b). We then asked whether a decrease in the synchrony of neurons switching between ON and OFF periods may explain the overall reduction in SWA, as predicted by modeling studies^25,26^.

We examined features of the LFP that reflect neuronal synchrony, including the LFP slope at OFF period initiation, the peak LFP amplitude within an OFF period, and the LFP slope at OFF period termination, all of which were reliably enhanced in the first 30 minutes of NREM recovery sleep compared to the 12-hour baseline. In both SOM+ and ACR mice, we found that all three measures were reduced on the optrode compared to the contralateral control probe (Fig. 2h-j), indicative of a reduction in neural synchrony after OFF period induction.

To examine synchrony at the level of neuronal firing (MUA), we computed the spike time tiling coefficient (STTC), a firing-rate-independent measure of neural synchrony^27^. On every channel-wise pair of MUA spike trains from each probe, the STTC measures the proportion of spikes from train-1 that have at least one spike from train-2 within a certain window (±5 ms), normalized by the chance level implied by each train’s own firing rate, thus providing a more rate-robust measure of spike train synchrony than traditional binned correlations (Fig. 2k). An example STTC matrix for the 1-hour NREM recovery period and the 1-hour NREM circadian matched period (the 1 hour of NREM sleep starting exactly 24 hours prior to the recovery period), for both the optrode and contralateral control probe is shown for a representative SOM+ animal in Fig. 2l. Again, we found a tight correspondence between the homotopic probes in the baseline period, followed by a dissociation during the recovery period (Fig. 2m,n, left). Across all subjects, the STTC was significantly reduced on the optrode relative to the contralateral control when averaging across all channel pairs for SOM+ (Fig. 2m) and ACR (Fig. 2n) mice. Thus, the decrease in SWA produced by the local induction of OFF periods is accompanied by a concomitant decrease of LFP and MUA markers of neuronal synchrony.

### An equivalent tonic reduction of firing rate cannot account for the effects of ON/OFF period induction

Is the reduction of SWA and cortical synchrony obtained by inducing OFF periods during wakefulness due to an overall reduction of firing rates, or does the sleep-like alternation of ON/OFF periods matter? If sleep pressure were to increase because of tonic activity during wakefulness, reduced firing during NREM sleep might be sufficient to relieve that pressure^28,29^, regardless of how reduced firing is achieved. To address this issue, we employed a third optogenetic approach paired with bilateral silicon probe recordings of homotopic cortical areas. We crossed CaMKIIα-Cre mice with mice expressing the inward chloride pump halorhodopsin (eNpHR3.0, Fig. 3a)^30^. Using these halorhodopsin mice (N = 6), we were able to tonically inhibit nearly all neurons in the unilateral cortical network during the final ∼30 minutes of SD (Fig. 3b-d).

**Fig. 3.**
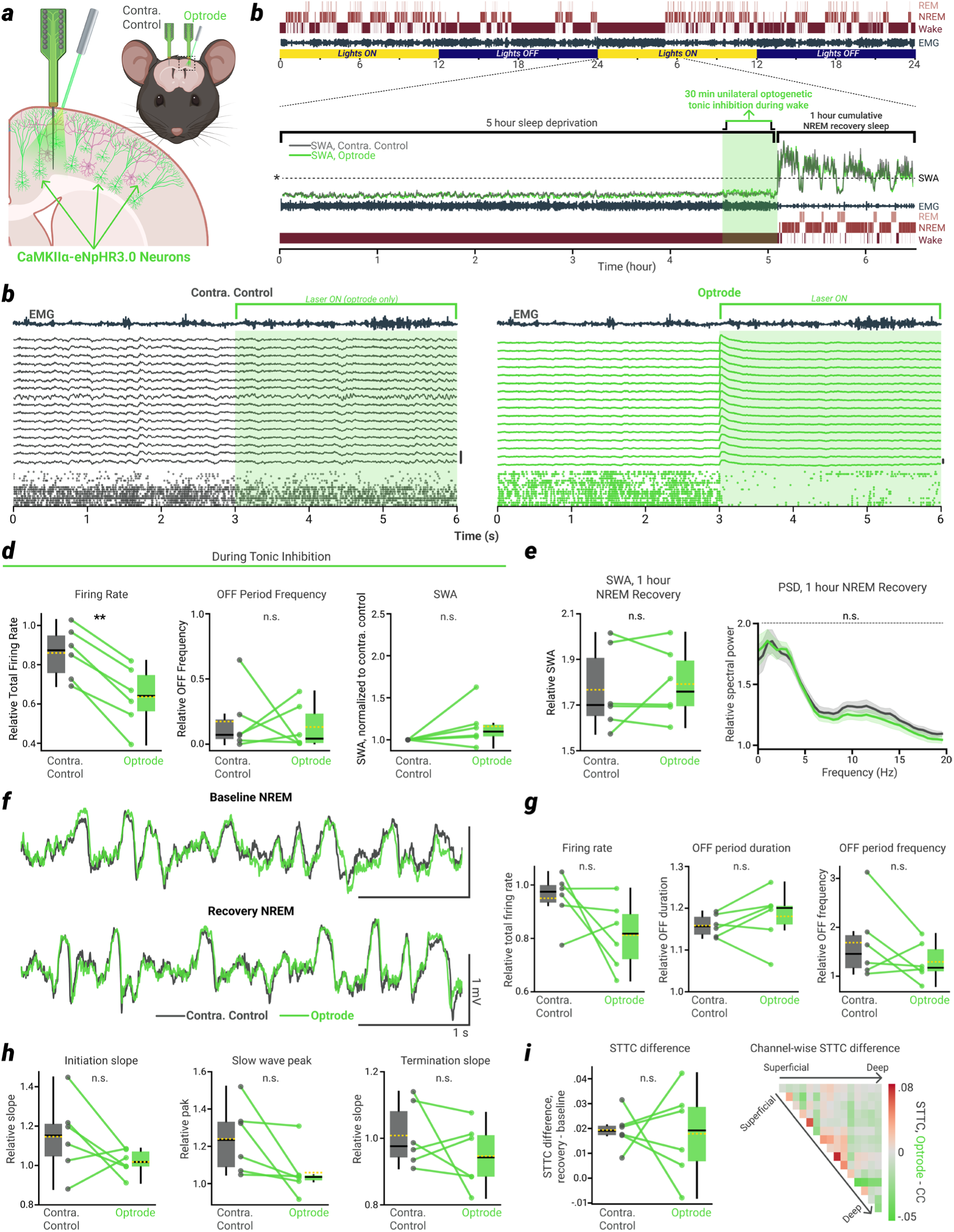
Tonic inhibition using halorhodopsin does not reduce SWA or synchrony in subsequent sleep. **a**, Halorhodopsin mice express *Natronomonas pharaonis*-derived halorhodopsin (eNpHR3.0) in CaMKIIα-producing neurons (mainly pyramidal neurons, green cells indicated by arrows). When eNpHR3.0 is optogenetically activated, all CaMKIIα neurons in the cortical network are inhibited via a light-driven chloride pump. **b**, Experiment overview. In all halorhodopsin mice, unilateral tonic inhibition of overall MUA firing rate was performed for the final ∼30 minutes of SD. All other aspects of the experimental design were identical to the OFF period induction experiments previously described (see materials and methods). *Dotted line indicates the 12 hour NREM baseline SWA mean. **c**, 6 seconds of raw data from a representative mouse showing tonic inhibition of MUA using continuous activation of eNpHR3.0 on the optrode, and not on the contralateral control probe. Scale bars at right of each plot represent 2 mV. **d**, Firing rate, OFF period frequency and SWA on the optrode relative to the contralateral control during tonic inhibition using eNpHR3.0. Firing rate is expressed relative to first hour of SD, OFF period frequency and SWA are calculated relative to the 12 hour NREM baseline mean, and SWA is expressed normalized to the contralateral control for visualization. **e**, SWA and PSD during the l hour NREM recovery period. **f**, Example NREM LFP traces in baseline and recovery sleep; note the tight correspondence between optrode and contralateral control LFP continues from baseline into recovery sleep. Shown is one matched layer 5 channel on each probe of a representative mouse (same mouse from **c**), with implant in M2. **g**, Total MUA firing rate, OFF period duration and OFF period frequency during the 1 hour NREM recovery period. **h**, Field potential slopes at OFF period initiation and termination, and slow wave peak values, during the 1 hour NREM recovery period. In **e**, **g**, and **h**, all data are expressed relative to the 12 hour NREM baseline mean. **i**, STTC difference (1 hour NREM recovery period mean – 12 hour NREM baseline mean). Right shows the data in channel-wise form, expressed as the difference between the optrode and the contralateral control. In all panels, *P<.05 | **P<.005 | ***P<.0005 | n.s. P>0.05.

Crucially, despite this large reduction of overall firing rate (roughly equivalent to the that seen during OFF period induction), NREM sleep-like ON/OFF periods were not triggered and SWA did not increase (Fig. 3d). Following this tonic firing rate reduction, NREM SWA on the optrode side was not significantly affected relative to the contralateral control probe (Fig. 3e). This was again evident in the raw NREM LFP traces, as the tight correlation between matched channels seen in the NREM baseline period continued into the NREM recovery period following SD (Fig. 3f). Across all other measures examined for the OFF period induction experiments, no significant effects were found on the optrode following the tonic inhibition of firing rate during wake. Thus, during the 1 hour NREM recovery period following tonic inhibition, OFF period duration and frequency, overall firing rate, LFP slope at OFF-initiation and OFF-termination, slow wave peak amplitude, and spike time synchrony showed no significant differences between optrode and contralateral control (Fig. 3g-i).

We further investigated the effects of tonic inhibition in a subset of SOM+ mice (N = 5). We performed a separate experiment where we tonically reduced firing rate via continuous sinusoidal stimulation of SOM-ChR2 neurons (Extended Data Fig.5a, b). Again, this resulted in an overall reduction of firing rate comparable to the OFF period induction protocol (Extended Data Fig. 5c), but few OFF periods (Extended Data Fig. 5d), and no significant reduction of SWA in subsequent NREM recovery sleep (Extended Data Fig. 5e). Critically, when directly comparing within the same mice the ability to reduce SWA in NREM recovery sleep, the induction of ON/OFF periods during wake was more effective than a tonic reduction of firing rate, in every mouse tested (Extended Data Fig. 5e, right).

These results provide converging evidence, across two in vivo optogenetic approaches, that the ON/OFF bistable activity patterns which characterize NREM sleep, but not mere reduction of firing, can release sleep pressure and reduce neural synchrony in a local cortical network even while the animal is awake and behaving. As such, all further experiments and analyses focus solely on the OFF period induction protocol, and the ability of ON/OFF activity patterns to fulfill key homeostatic sleep functions when induced during wake.

### Induction of OFF periods during wakefulness leads to a reduction of molecular markers of synaptic strength

Modeling studies also predict that the homeostatic decline of SWA and synchrony during sleep (together with the decline of slow wave amplitude and slope, and overall firing rate, Extended Data Fig. 4c) can be accounted for by a net reduction of excitatory synaptic strength in cortical networks^25,26^. Such a reduction has been confirmed experimentally using electrophysiology^31,32^, ultrastructural measurements of synaptic size and density^33,34^, as well as molecular markers^9,10,31^. We therefore took advantage of the same molecular markers of synaptic strength to determine whether the induction of OFF periods during wakefulness can have similar effects as sleep on cortical synaptoneurosomes.

We implanted SOM+, ACR, and wildtype control mice (N = 8/group) with EEGs over parietal cortex. On one side of the brain, 4 optic ferrules were implanted within a 3 mm diameter circle over parietal cortex, with each ferrule 1 mm from the ‘opto-EEG’ (Fig. 4a), thus enabling us to induce NREM sleep-like ON/OFF bistable activity throughout a large area of unilateral parietal cortex. These mice underwent the same SD procedure as those implanted with silicon probes, except following the 30 minutes of unilateral OFF period induction at the end of SD, mice were immediately sacrificed and were not allowed to sleep. A mirrored 3 mm diameter circle of cortex was collected from each side of the brain, centered over each of the bilateral EEGs (Fig. 4a, left). Using this tissue, synaptoneurosomes were extracted and two established markers of excitatory synaptic strength^35^ were measured – the amount of cortical synaptic expression of AMPA (alpha-amino-3-hydroxy-5-methyl-4-isoxazole propionic acid) receptors containing the GluA1 subunit, and the amount of phosphorylation of GluA1 at serine 845 (pGluA1(845)).

**Fig. 4.**
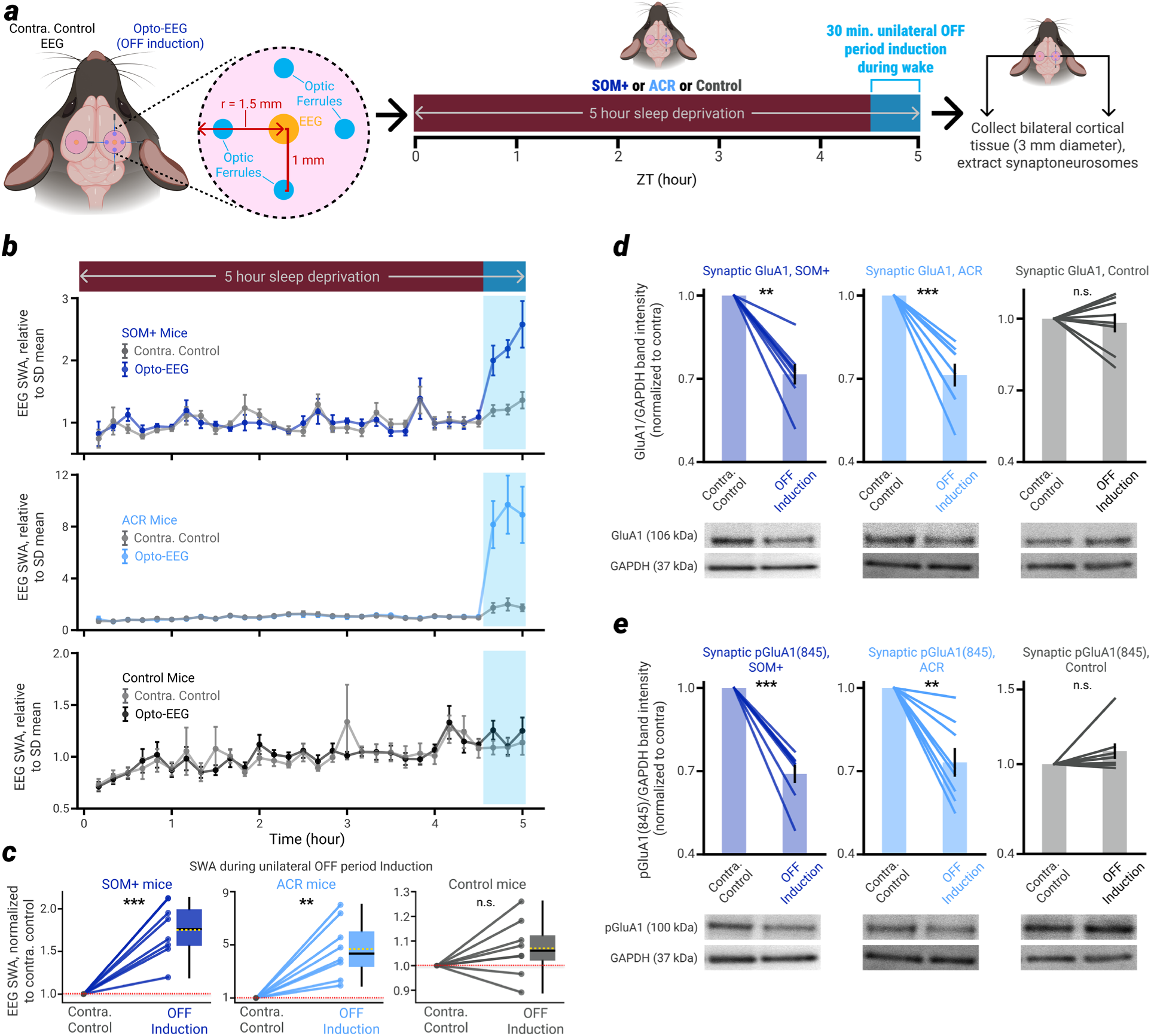
OFF period induction during wake reduces molecular markers of cortical excitatory synaptic strength **a**, Experiment overview. SOM+, ACR, and control mice are implanted with bilateral EEGs over parietal cortex and 4 optic ferrules on one side. OFF period induction in one half of the brain occurs at the end of SD (final 30 minutes), followed by brain collection. A 3mm diameter circle of cortical tissue from each half of the brain (pink circular region) was used for synaptoneurosome extraction. **b**, SWA during SD in the opto-EEG and the contralateral control EEG for SOM+, ACR, and control mice. **c**, Average EEG SWA during OFF period induction, normalized to contralateral control EEG. **d**, Synaptic levels of GluA1 for both hemispheres in SOM+, ACR, and control mice. Representative immunoblots are shown for each experimental group. Data were normalized by the intensity of the GAPDH band and expressed as means ± S.E.M. **e**, As in **d**, but for synaptic levels of phosphoGluA1. Brightness and contrast were adjusted for all blot snippets in this figure, solely for space and clarity considerations. In all panels, *P<.05 | **P<.005 | ***P<.0005 | n.s. P>.05.

In both SOM+ and ACR mice, EEG recordings confirmed that the OFF period induction was accompanied by a large increase in SWA in the corresponding hemisphere (Fig. 4b,c). In both SOM+ and ACR mice (but not in controls), the cortical hemisphere that underwent OFF period induction also had significantly lower synaptic levels of GluA1-containing AMPA receptors (Fig. 4d) and their phosphorylation at serine 845 (Fig. 4e, Extended Data Fig. 6). In both direction and magnitude, these changes mirror those seen after natural NREM sleep^10,31^.

Since in these experiments mice were not allowed any sleep following SD, the changes in AMPA receptors must have been triggered by the forced induction of NREM sleep-like ON/OFF activity patterns during wakefulness.

### Induction of OFF periods during SD recovers memory performance in a novel floor texture recognition task

There is substantial evidence that sleep plays a critical role in learning and memory consolidation^8^. Moreover, at least some of these benefits of sleep may be mediated by sleep-dependent synaptic down-selection. For example, after learning a complex wheel task, adult mice that are allowed to sleep immediately following learning perform better in the next training session than mice that are sleep deprived, and the weakening of non-task-engaged excitatory synapses is correlated with that improvement (revealed through two-photon microscopic imaging of SEP (super ecliptic pHluorin)-GluA1 expression^36^). This raises the question of whether inducing sleep-like ON/OFF periods during wakefulness can provide a memory or learning benefit similar to that seen after sleep.

To test this possibility, we employed a floor texture recognition (FTR) task that is known to benefit from sleep and involves neuronal activity localized to secondary motor cortex (M2) and primary sensory cortex (S1)^17^. We focused on SOM+ mice because of the involvement of somatostatin-positive interneurons in the generation of NREM sleep slow waves in natural conditions^23^ and implanted them with bilateral optic ferrules over M2 and S1 (Fig. 5a). Mice were exposed to an arena with identical floor textures on either side of the chamber for 15 minutes (‘acquisition’ period). Following acquisition, mice were returned to their home cage and either allowed to sleep freely (sleep group, N = 9), sleep deprived for 1 hour (SD group, N = 13), or sleep deprived for 1 hour with concurrent induction of NREM sleep-like OFF periods (SD + OFF induction group, N = 8). Mice were recorded in their home cages, and we verified the presence of sleep in the group of mice immediately allowed to return to their cage and sleep (Extended Data Fig. 7). 24 hours following acquisition mice were placed back into the arena, where now half of the arena contained a floor with a novel texture (‘recall’ period, Fig. 5b). We then used a neural network trained to recognize the mouse’s position within the arena to quantify the amount of time the mouse spent on each side of the arena within the first 5 minutes of the recall period^37^. Replicating previous findings^17^, we found that mice allowed to sleep immediately following acquisition performed better than mice sleep deprived for 1 hour following acquisition (Fig. 5c,d). Moreover, in SOM+ mice that underwent bilateral induction of NREM sleep-like OFF periods during 1 hour of SD following acquisition, performance was rescued back to the level of the mice allowed to sleep (Fig. 5c,d). Thus, inducing sleep-like ON/OFF periods during wakefulness can benefit memory consolidation similar to sleep.

**Fig. 5.**
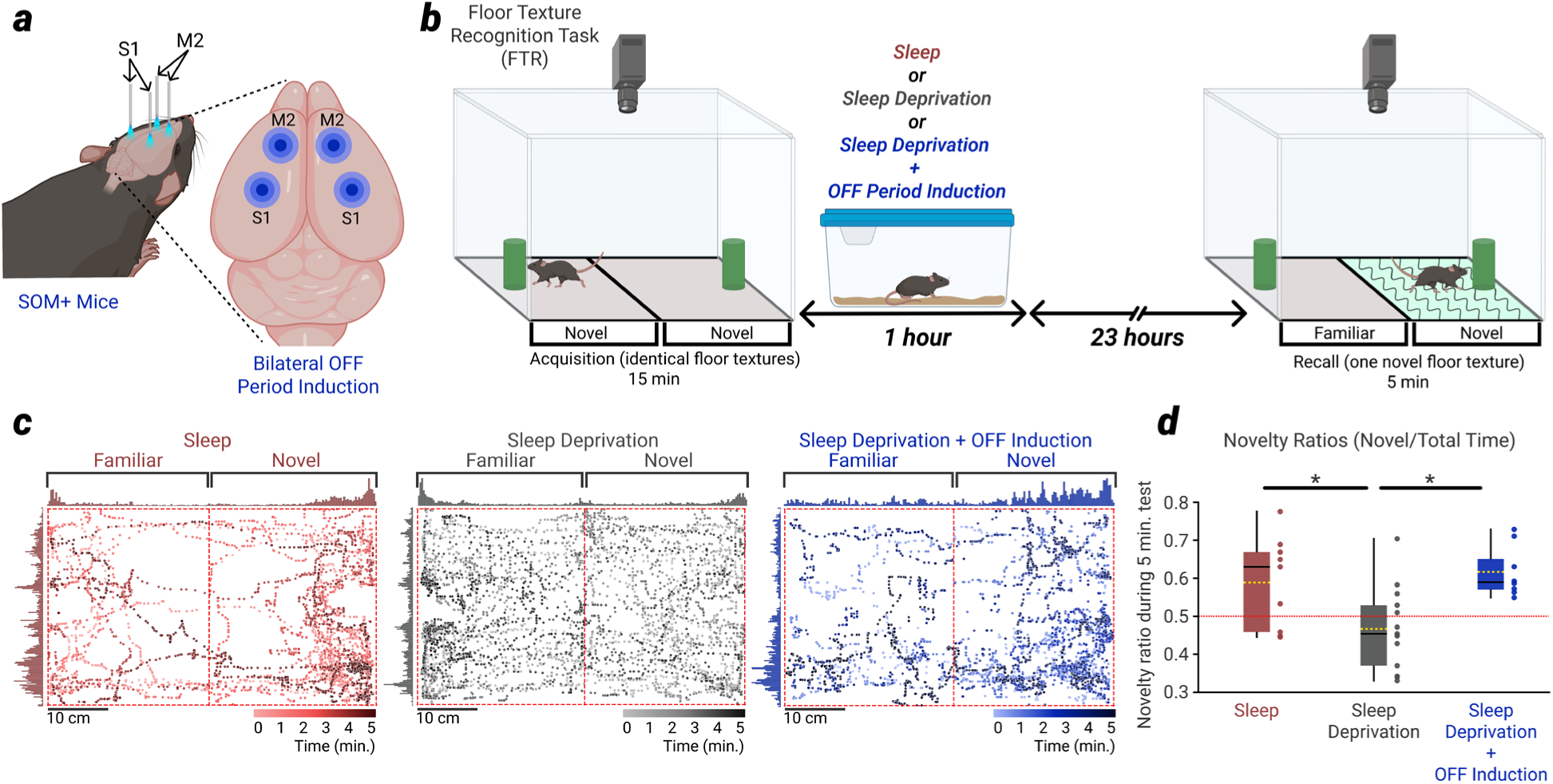
Bilateral OFF induction during SD recovers memory performance in a novel floor texture recognition task **a**, Implant schematic. Optic ferrules are implanted bilaterally over M2 and S1 cortices. **b**, Following acquisition of the floor texture recognition task (FTR), mice return to the home cage where they are either allowed to sleep, sleep deprived for 1 hour, or sleep deprived while OFF periods are induced for 1 hour. The next day mice are tested on FTR. **c**, A time-resolved plot showing the neural network’s labelling of the mouse’s position within the arena over the 5-minute recall period for a representative mouse from the sleep, SD, and SD+OFF induction groups. **d**, Performance (ratio of time spent on novel side relative to familiar side) for each of the three groups.

## Discussion

We show here that sleep-like neuronal OFF periods can be reliably induced in local cortical networks while mice are awake and behaving. OFF periods induced during sustained wakefulness locally reduce sleep pressure, as assessed by a decline of SWA and neuronal synchrony, compared to the contralateral homotopic cortex. Sleep pressure is relieved if OFF periods alternate with ON periods every few hundreds of milliseconds, as it happens in NREM sleep, while an equivalent overall reduction of neuronal firing rate obtained through tonic inhibition is not effective. The induction of sleep-like OFF periods during wakefulness also leads to a decrease in excitatory synaptic strength, as assessed by a reduced expression and phosphorylation of GluA1-containing AMPA receptors, similar to what happens after NREM sleep. Finally, inducing OFF periods bilaterally over motor and sensory cortices during waking after learning restores performance in a recognition task, similar to what sleep does. Thus, local OFF periods occurring during wakefulness can offer key benefits normally provided by sleep.

During NREM sleep, the induction and maintenance of the characteristic bistability between synchronous ON/OFF periods is enabled by the low level of activity of all major arousal systems, which leads to neuronal disfacilitation. Blocking noradrenergic transmission during wakefulness is sufficient to trigger local OFF states in the barrel cortex^38^, consistent with in vitro evidence that noradrenaline and acetylcholine strongly oppose cortical ON/OFF bistability^39^. Active inhibition by cortical SOM+ interneurons also plays a role in promoting bistability^23,40,41^. SOM+ interneurons are activated by strong synchronous firing of pyramidal neurons^42,43^ through progressively facilitating synapses^44–47^. When enough SOM+ interneurons are recruited, they act as “master regulators” of cortical excitability by inhibiting all other cell types but themselves^48–51^. As shown here, inhibition of pyramidal neurons via SOM+ interneurons is sufficient to induce local OFF periods during waking for as long as we tried (30-60 minutes). Direct inhibition via stGtACR1 can also produce OFF periods, though the mechanisms are likely less physiological, as indicated by the opposite polarity of the field potential compared to spontaneously occurring and SOM+ induced OFF periods (Fig. 1h).

The occurrence of OFF periods during NREM sleep underlies SWA^15,52^. In fact, increased SWA is associated with more synchronous and longer OFF periods^16^. SWA is a reliable maker of sleep pressure in both rodents and humans^53,54^. There is also abundant evidence that the level of SWA, as well as the amplitude and slope of individual slow waves, reflect changes in net cortical synaptic strength, increasing with the duration of wakefulness and decreasing homeostatically during sleep^5,31,55,56^, in line with modeling studies^25,26^. An intriguing possibility is that SOM+ interneurons may be involved in mediating the increased occurrence, duration, and synchrony of OFF periods associated with increased sleep pressure. Synaptic activation of NMDA receptors on the dendrites of cortical pyramidal neurons leads to long-term potentiation of the inhibitory synapses established by SOM+ interneurons, suggesting a mechanism through which wake-induced potentiation of excitatory cortical synapses could lead to stronger SOM+-mediated dendritic inhibition and increased propensity to enter an OFF state^57^.

It has been suggested that SWA and associated OFF periods do not simply reflect sleep pressure as a consequence of net synaptic potentiation, but may be causally involved in its homeostatic regulation by enacting synaptic renormalization^5,58^. In vitro experiments in rodent^59–61^ and human^62^ cortex show that bistable, ON/OFF conditions are typically associated with synaptic depression, although the underlying mechanisms remain unclear. In vivo experiments under urethane anesthesia are consistent with this conclusion^63^, but the evidence in naturally sleeping animals is limited and indirect^64^. The present results demonstrate in vivo that sleep-like ON/OFF alternations lead to the weakening of excitatory glutamatergic synapses. This is directly confirmed by the reduced levels of GluA1-containing AMPA receptors and their phosphorylation, and indirectly by the reduced amplitude and synchrony of ON/OFF periods and associated slow waves during subsequent sleep. The effect on synchrony was not obtained by an equivalent reduction in firing rate induced by tonic inhibition, suggesting that repeated sequences of ON/OFF transitions may be critical for synaptic depression.

Remarkably, OFF period induction was effective in relieving local sleep pressure when performed in awake animals. Assuming that the overall levels of noradrenaline and acetylcholine were those typical of wakefulness (high), rather than of NREM sleep (low), our results would suggest that low levels of neuromodulators may be permissive for enabling bistable activity, but not necessary for producing synaptic renormalization, as long as ON/OFF periods can be triggered through other means. In vitro experiments have indicated that low levels of noradrenaline coupled with low levels of brain-derived neurotrophic factor may be conducive to synaptic depression and the removal of AMPA receptors from the synaptic membrane^65–67^. It will be important to establish whether the local induction of OFF periods may indirectly affect the local level of neuromodulators. Previous experiments in mice found that the removal of AMPA receptors depends on the activation of constitutive, ligand-independent mGluR1/5 signaling following the accumulation of Homer1a in the dendritic spines, which requires the drop in noradrenaline levels as typically observed at sleep onset, or the build-up of adenosine in the course of sleep deprivation^65^. In our experiments, adenosine levels likely increased on both sides of the cortex after 4.5 hours of sleep deprivation, but synaptic strength decreased on the side with OFF induction. The weakening of cortical synapses may thus require the occurrence of activity bursts, perhaps tied to the activation of mGluR1/5 signaling, as observed for cerebellar and hippocampal synapses^68^.

In previous work, we showed that extended periods of sustained wakefulness, especially if coupled with intense learning, lead to the spontaneous occurrence of ON/OFF patterns of activity, which we termed “local sleep”^69^. Local sleep has been demonstrated in both rodents^69^ and humans^70^. Its spontaneous occurrence is sporadic and can disrupt performance if it occurs in the wrong brain region at the wrong time^69^. The present findings raise the question whether local sleep during prolonged wakefulness, if sustained for many minutes, may help to control local synaptic strength and protect local circuits from negative consequences on cellular processes, thus subserving some of the functions of sleep. A systematic induction of ON/OFF transitions over specific networks in an otherwise awake brain, may be much more effective in restoring homeostasis and performance than sporadic OFF periods. Species such as bottlenose dolphins, beluga whales, mallard ducks, and frigate birds evolved unihemispheric NREM sleep to enable its restorative function while maintaining vigilance with the other hemisphere^71^. Conceivably, technologies capable of inducing ON/OFF periods on demand in select brain regions may provide some of the benefits of sleep without paying the full price of sensory and motor disconnection. On the other hand, even if sleep can be induced and regulated locally, the overall disconnection enforced by sleep is likely necessary to ensure that synaptic renormalization and memory consolidation can take place not just for local circuits, but at the systems level, enabling the spontaneous activation of brain-wide circuits^7^.

## Methods

### Animal husbandry

All surgical and experimental protocols described in this study were approved by the University of Wisconsin Institutional Animal Care and Use Committee (IACUC). Mice were housed on a 12:12 light:dark cycle, room temperature was maintained at 23°C ± 1°C, and 50% humidity. All experiments used adult mice (>8 weeks at time of surgery); male and female mice were used in roughly equal proportions in all experiments, based on availability at the time of surgery.

The ACR mouse line was generated by crossing R26-LNL-GtACR1-Fred-Kv2.1 male mice (Jackson Labs #033089) with B6.Cg-Tg(Camk2a-cre)T29-1Stl/J female mice (Jackson Labs #005359). Heterozygous offspring were used as wildtype controls. The SOM+ mouse line was generated by crossing Ai32(RCL-ChR2(H134R)/EYFP) male mice (Jackson Labs #024109) with Sst-IRES-Cre female mice (Jackson Labs #013044). All resulting offspring were homozygous and thus were not genotyped. The Halorhodopsin mouse line was generated by crossing Ai39(RCL-eNpHR3.0/EYFP) male mice (Jackson Labs #014539) with B6.Cg-Tg(Camk2a-cre)T29-1Stl/J female mice (Jackson Labs #005359). All resulting offspring were homozygous and thus were not genotyped.

## Surgery

### Preparation of mice for silicon probe electrophysiology experiments

All silicon probes used in this study consisted of 16 channels in a linear array, with 50 µm channel spacing (NeuroNexus Technologies; A1×16-3mm-50-177-CM16LP). The ‘optrodes’ referred to in these experiments were manually constructed by physically attaching an optic ferrule (Doric Lenses Inc., MFC_200/240-0.22_8mm_MF2.5_DFL) to a silicon probe. The desired target location in the brain for the electrodes on the silicon probes was planned, and the optic ferrule was cemented to the probe such that the tip of the ferrule would end up just at the surface of the brain when the probe was inserted to its final depth (i.e. the ferrule did not penetrate into the brain). The ferrule was positioned such that its tip was ∼150-200 µm from the shank of the probe, with a 12-15 degree angle relative to the shank.

Mice were implanted with bilateral silicon probes in homotopic cortical areas (i.e. at the same anterior-posterior (AP) and dorsal-ventral (DV) coordinates, and at mirrored medial-lateral (ML) coordinates) of either the frontal or parietal cortex. The stereotaxic coordinates relative to bregma (the DV axis is relative to the brain’s surface) for the implant locations were as follows: 1) for frontal animals with an implant in secondary motor area (M2): AP +2.4 mm, ML ±1.6 mm, DV –1.15, θ: 3°; 2) for parietal animals with an implant in primary somatosensory cortex (S1): AP –1.4 mm, ML ±2.0 mm, DV –0.97 mm, θ: 0°; 3) for parietal animals with an implant in posterior parietal cortex (PPC): AP –2.0 mm, ML ±2.15 mm, DV –0.92 mm, θ: 4°.

Before the surgery, mice were given one dose of ceftriaxone (100 mg/kg) and one dose of dexamethasone (2.5 mg/kg). They were allowed ad libitum oral carprofen (Bio-Serv) for one day before and 3-6 days following surgery. Mice were anesthetized using 1-2% isoflurane and body temperature was maintained using an electronic heating pad. Mice were fixed onto a stereotaxic rig (Kopf Instruments, model 942), and the skull was exposed. A circular craniotomy 1 mm in diameter was drilled over the target location for each probe. One stereotaxic arm was then used to position each probe at the target location over the craniotomy, and the probes were slowly lowered to the required depth in the brain. The craniotomies were sealed with a biologically safe silicone polymer (‘Kwik-Sil’; World Precision Instruments), and the probes were cemented to the skull with dental cement (‘C&B Metabond’; Parkell, Inc.). Reference and ground screws were placed over the cerebellum. Stainless steel EMGs were implanted into the nuchal muscles at the end of the surgery.

All mice were given at least 1 week of full undisturbed recovery following surgery. Once mice had recovered, they were briefly anesthetized with 1-2% isoflurane and connected to the electrophysiology acquisition system and the implanted ferrule(s) was connected to a patch cord (Doric Lenses, Inc.) for delivery of light in optogenetic experiments.

### Preparation of mice for quantitative immunoblotting experiments

For mice used for quantitative immunoblotting experiments, EEGs, optic ferrules, and nuchal EMGs were implanted. First, two 0.7 mm burr holes were drilled and two stainless steel screw EEGs were implanted over parietal cortex (AP −2.0, ML ±2.5). Then, four 0.5 mm burr holes were drilled, each 1 mm away from one of the EEGs on only one side of the brain (see Fig. 4). Optic ferrules were slowly lowered into the hole until they reached the brain surface, and the hole was sealed with a clear silicone polymer (Kwik-Sil) before all EEGs and ferrules were further cemented to the skull. Reference and ground screws were implanted over cerebellum. Surgical procedures were otherwise identical to those described above for mice receiving bilateral silicon probe implants.

### Preparation of mice for novel floor texture recognition experiments

For mice used in tests of novel floor texture recognition, four optic ferrules were implanted. Two ferrules were implanted bilaterally over M2 (AP +2.20, ML ±0.65), and two were implanted bilaterally over S1 (AP –0.73, ML ±1.95). All other surgical procedures were otherwise identical to those described in the above two sections.

### Electrophysiology acquisition and analysis

#### Electrophysiology data acquisition

For the acquisition of EEG and EMG signals, copper hookup wire was used to connect the animal to the electrophysiology acquisition system. Raw EEG and EMG data were acquired and stored at 1,017 Hz. Silicon probes were connected to the electrophysiology acquisition system via low profile head stages (LP16CH-Z, Tucker-Davis Technologies). Raw data from the silicon probes were acquired and stored at 24.414 kHz. All electrophysiology data were acquired using a battery-powered amplifier (PZ5, Tucker-Davis Technologies) in conjunction with a signal processor (RZ2, Tucker-Davis Technologies) and standard acquisition software (Synapse neurophysiology suite, Tucker-Davis Technologies).

#### Electrophysiology data processing

Generally, all electrophysiology analysis was done using custom code written in python. Code required to reproduce statistical tests and plots in the paper (as well as all statistical source data) is available at the following repository: (https://github.com/CSC-UW/OFF_PERIOD_INDUCTION_MATERIALS). For MUA-based analyses, raw 24.4kHz data acquired from the silicon probes were bandpass filtered between 300Hz and 12kHz, and a global common median reference was applied. Noise levels on the filtered data were estimated using median absolute deviations, and a spike threshold was set at four times the estimated noise level. For every channel, peaks were detected when the threshold was crossed, and spike times were recorded. Where firing rates are reported, firing rate was computed in 2 second windows across the duration of the recording. All MUA preprocessing and detection were done using standard routines from the spikeinterface python package^72^.

For LFP-based analyses, raw 24.4kHz data were low pass filtered using a finite impulse response filter and decimated to 400Hz. Raw EEG and EMG data were acquired at 1017 Hz and not decimated before analysis. After all data processing, bad channels were marked manually, interpolated where possible (via a simple spatial interpolation using the voltages on neighboring channels^72^), and excluded from a given analysis if interpolation could not resolve data quality or stability problems, though this was rare. Throughout the paper, when silicon probe data or analyses derived from those data are presented at the level of comparisons between animals or probes (e.g. time-frequency analyses, firing rate analyses, synchrony analyses, etc.), data always reflect the average of all viable channels across the probe, unless otherwise noted.

#### Time frequency analyses

Time frequency analyses were performed on LFP or EEG data following the preprocessing steps outlined above. Power spectral density (PSD) was estimated with a short-time Fourier transform using 4s Slepian (DPSS) windows with time–half-bandwidth product NW=4 and 50% overlap (2s hop). SWA was computed by integrating the spectral power in the delta band (0.5 – 4 Hz). SWA was expressed relative to the NREM mean of the 12-hour light-period baseline value.

#### OFF period detection and synchrony analyses

OFF periods were first detected at the single-channel level using a given channel’s MUA spike train. At the single channel level, an ON period was defined as any period of continuous spiking activity with no interspike intervals greater than or equal to 50ms. OFF periods were defined as any period of total silence lasting at least 50ms, with an ON period at least 30 ms in duration on either side (before or after) of the OFF period (i.e. an OFF period cannot be surrounded by two single-spike “ON periods”). Global OFF periods were then detected as periods where most channels (12 or more out of 16) were synchronously in an OFF period for between 50 and 400 ms. Global ON periods were defined as those periods where less than 12 channels were synchronously OFF for between 50 and 2000 ms. Unless otherwise noted, all OFF period analyses presented throughout the paper refer to the global OFF period definition. In rare cases with very low MUA on a number of channels, the definition of OFF period used above, which requires OFF periods to be preceded and followed by more than one spike, resulted in an extremely low OFF period detection rate. In these cases, OFF periods were defined as a period of zero MUA spiking across all channels lasting at least 50 ms, as in reference^16^, and the same definition and detection method were always applied to both probes for a given subject (optrode and contralateral control).

To detect OFF period initiation slopes, trace snippets for each channel were extracted and aligned relative to all OFF period start times within a given condition (e.g. 1-hour NREM recovery sleep). Then, an average waveform was calculated for all channels across all OFF periods in the condition. The first order derivative of these traces was computed after applying a Savitsky-Golay filter, and the maximal value of the first order derivative within ±10 samples of the OFF period start sample was taken as the maximal initiation slope for that channel. The same procedure was followed for OFF period termination slope, except traces were aligned to the OFF period end, rather than the start. To detect the OFF period-associated slow wave peak values, trace snippets for each channel were extracted and aligned to the midpoint of the OFF period.

Average waveforms were calculated for all channels across all OFF periods, and the peak value of the field potential within ±20 samples of the OFF period midpoint sample was recorded.

Spike time tiling coefficients (STTC) were computed as described in^27^. Briefly, for a given pair of spike trains *A* and *B*, we computed the proportion of spikes in each train that had at least one coincident partner spike in the other train within a ±5 ms window (TA, TB), and the fraction of the total recording time that already lay within ±5 ms of any spike in the same train (PA, PB, i.e. the union of [*t* – Δ*t*, *t* + Δ*t*] around spikes of A or B, respectively), which were then combined to yield the symmetric and rate-corrected metric below:

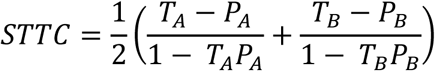

STTC was computed for every unique channel pair on a per-epoch basis (i.e. for each continuous bout within a condition) and all epochs were averaged across the entire condition. STTC values were Fisher-Z transformed (*z*=atanh(*r*)) after clipping *r* between [-1+ ε, 1-ε] where ε=1×10^-6^) before computing condition means or differences in the coefficient (e.g. between baseline and recovery sleep, or between optrode and contralateral control) or running statistical tests.

#### Sleep scoring

Arousal states (Wake, NREM sleep, REM sleep, or intermediate state in the transition to REM sleep) were manually assigned using custom software written in python (https://github.com/CSC-UW/sleepscore), based on a fork of visbrain-sleep^73^. When EEGs were available, scorers used all EEG channels to make state decisions. When only LFPs from silicon probes were available, experiments used a fraction of channels (∼1/4) spanning the probes to make state decisions. Ambiguous or unclear states were marked as ‘Unsure’, though this constituted a very small minority of all labelled data (<0.5%). Artifact was manually marked where obvious and feasible, but because of the variance across channels and probes, it was largely not possible to comprehensively label artifactual periods manually. As such, bad channels were individually interpolated on an as-needed basis, and artifactual channels or epochs were further excluded on a per-experiment basis as needed. Scorers labelling data were blinded to animal and experiment identity.

Unless otherwise noted, the ‘baseline’ condition refers to all periods scored as NREM sleep during the 12-hour light period on the day before the experiment. The ‘recovery sleep’ condition refers to the first 1 hour of cumulative NREM sleep following SD. The ‘circadian-matched baseline’ condition refers to the first 1 hour of cumulative NREM sleep beginning exactly 24 hours before the exact start of the recovery sleep condition.

#### Sleep deprivation and optogenetics

The day before all electrophysiology experiments described in this paper, a 24-hour baseline recording was acquired, starting at light onset on the day before the experiment (i.e. 24 hours before the experiment begins). The experiment consisted of a 5-hour sleep deprivation (SD) by exposure to novel objects, beginning at light onset on the day of the experiment. After placing a novel object in the home cage, the mouse explored the object for a period of time; when the mouse became disinterested in the object, stopped exploring, or looked as if it was preparing for sleep (nesting, grooming, etc.), the old object(s) was removed, and a new object(s) was placed into the cage. This process was repeated for the entirety of the 5-hour SD.

During the final 30 minutes of the SD, OFF periods were selectively and optogenetically induced in the cortical network recorded by the optrode. Importantly, the SD procedure continued throughout the entirety of this 30 minute OFF period induction period. The pattern of OFF period induction was planned to start with longer induced OFF periods, and progressively move to shorter durations, in order to mimic the natural decline in OFF period duration seen after SD in NREM sleep^16^. Indeed, the temporal protocol we used typically produced a rise and then subsequent decline in SWA, as is naturally seen in NREM sleep (Extended Data Fig. 2a). There were 30-45 second breaks in between bouts of OFF induction, to limit the duration of continuous light exposure and to mimic the wake-like brain activity seen during REM sleep or during brief arousals. The OFF period induction pattern consisted of the following stages: 1) Stage 1: Pulses delivered at 1 Hz, with a laser-on time of 180 ms, for 10 minutes total. Laser-off break: 45 seconds; 2) Stage 2: Pulses delivered at 2 Hz, with a laser-on time of 140 ms, for 10 minutes total. Laser-off break: 45 seconds; 3) Stage 3: Pulses delivered at 3 Hz, with a laser-on time of 100 ms, for 5 minutes total. Laser-off break: 45 seconds; 4) SD ends – at this point the formal SD stopped, all objects were removed from the cage, and the mouse was free to sleep at any time; 5) Stage 4: Pulses delivered at 3 Hz, with a laser-on time of 80 ms, for 5 minutes total.

After each 5-minute bout of stage-4, the laser was turned off for a 30 second break, after which another bout began. This stage 4 continued until the mouse actually fell asleep. In short, the SD formally ended (i.e. efforts to keep the mouse awake ended) 5 hours after light onset, but OFF period induction continued until the mouse actually began to sleep, at which point it was immediately and manually terminated. Thus, the actual duration of OFF period induction varied slightly across mice, typically 30 to 45 minutes. The recording continued until light onset of the following day.

For OFF period induction in the synaptoneurosome experiments, the induction protocol was largely identical to that used in silicon probe electrophysiology experiments, and OFF periods were again only induced during the final 30 minutes of the 5 hour SD. Specifically: 1) Stage 1: Pulses delivered at 1 Hz, with a laser-on time of 180 ms, for 10 minutes total. Laser-off break: 45 seconds; 2) Stage 2: Pulses delivered at 2 Hz, with a laser-on time of 140 ms, for 10 minutes total. Laser-off break: 45 seconds; 3) Stage 3: Pulses delivered at 3 Hz, with a laser-on time of 100 ms, for 5 minutes total. Laser-off break: 45 seconds; 4) Stage 4: Pulses delivered at 3 Hz, with a laser-on time of 80 ms, for 5 minutes total; laser-off break: 30 seconds. Critically, in these experiments the SD did not end, and mice were not allowed to sleep. Immediately following stage 4, the mouse was manually disconnected from optic patch cables and EEG/EMG tethers (pulses continued at 3 Hz/80ms as in stage 4 for this brief ∼1 min disconnection period), briefly anesthetized, and sacrificed for synaptoneurosome collection.

A 470nm laser (OEM Laser Systems DPSSL Driver) was used for all OFF induction experiments and any experiments involving either ACR or SOM+ mice. Power at the ferrule tip was measured using a photodiode power sensor (Thorlabs S120C) before optrode construction and surgical implantation. Power at the ferrule tip varied between mice and was adjusted within experiments at the discretion of the experimenter to achieve effective OFF period induction and/or tonic inhibition. These adjustments were limited to a small range, and the power in any optogenetic experiment (excluding the Halorhodopsin experiments) never exceeded ∼1.5mW. For OFF period induction in ACR mice, power at the ferrule tip was generally between 250 – 550 µW and was never greater than ∼700 µW. For OFF period induction in SOM+ mice, power at the ferrule tip was generally between 350 – 700 µW and was never greater than ∼1 mW. For OFF period induction in the synaptoneurosome experiments, slightly higher powers were used to maximize the diffusion of light and the area of affected tissue. Power at the ferrule tip was generally between 0.6 – 1.2 mW, and was never greater than 1.5mW.

For tonic inhibition experiments, the temporal pattern of light delivery was designed to match relatively closely the total MUA firing rate reduction observed in the OFF induction experiments, but without inducing many synchronized ON/OFF periods as in the OFF period induction experiments. Therefore, beginning 30 minutes before the end of SD, rather than inducing OFF periods as above, we delivered light continuously for 20 seconds, and then turned the light off for 60 seconds. This pattern was repeated for ∼30 minutes while the SD continued, as in the OFF induction experiments. After 30 minutes, the SD formally ended and the mouse was free to sleep, and this tonic inhibition protocol continued until the moment the mouse first fell asleep, at which point the inhibition was immediately and manually terminated and the mouse was left undisturbed, as in the OFF induction experiments.

For the experiments involving tonic inhibition using SOM+ mice, an analog voltage signal with a sinusoidal profile was used to drive the laser, at a frequency of either 0.5 Hz (1 mouse) or 40 Hz (4 mice). In all experiments, laser power at the ferrule tip was between ∼200uW and ∼700uW, and the distance between any given consecutive peak and trough was always between ∼100-200 µW. For the experiments involving tonic inhibition using Halorhodopsin mice, a 561 nm laser was used (OEM Laser Systems DPSSL Driver); the power at the ferrule tip was never more than 12mW and was typically in the range of 3-6 mW. The same experimental design and temporal pattern of light delivery was used as in the SOM+ tonic inhibition experiments, with two differences: first, the laser was driven digitally according to a TTL pulse, so when the laser was on, it was on at a consistent power rather than varying with the sinusoidal profile used in the SOM+ tonic inhibition. Second, in half of these mice, we used a 3-hour SD with inhibition during the final 30 minutes, rather than a 5-hour SD. This was done because of the known effects of sleep/wake history on intracellular chloride^74^ and our own experience suggesting it was in some mice easier to achieve effective inhibition at lower irradiance levels when sleep pressure was not as high as after 5 hours of continuous SD. These parameters and variations are largely taken from and in line with other published literature using these tools^74,75^.

### Quantification of synaptic GluA1 levels

#### Synaptoneurosome preparation

Following the 30 minutes of OFF period induction at the end of SD, mice were immediately deeply anesthetized with 1-3% isoflurane and decapitated. The left and right cortex was isolated, and a circular sample of diameter 3 mm was extracted from both hemispheres, centered over the EEG electrode, with the goal of collecting the majority of the affected tissue. Crude synaptosomes were prepared from fresh cortical tissues using Syn-PER synaptic protein extraction reagent (Thermo Fisher; 87793) supplemented with protease inhibitor cocktail (Halt Protease Inhibitor Cocktail, EDTA-Free (100X); 87785) following manufacturer’s instructions. Briefly, the samples were homogenized using a 2mL Dounce tissue grinder with 10 up-and-down even strokes. The homogenate was first centrifuged at 1200 × g for 10 minutes to eliminate cell debris, followed by a second centrifugation at 15,000 × g for 20 minutes. The pellets, containing synaptosomes, were gently resuspended in the respective amount of Syn-PER synaptic protein extraction reagent. Following extraction the samples were stored at −80 °C until used for western blot analyses.

#### Quantitative immunoblotting

For quantification of GluA1 and pGluA1 levels in the synaptoneurosome, we followed a protocol previously described in reference^10^. Protein concentration was measured using a Qubit 2.0 Fluorometer (Invitrogen, Life Technologies; Q32866). Aliquots were thawed on ice and denatured in a buffer containing Coomassie G250 and phenol red (Invitrogen™ NuPAGE) at 70°C for 10 min. Then, protein samples (10 μg) were loaded and separated electrophoretically using a 1.5 mm thick 4%–12% Bis-Tris gel together with NuPAGE MOPS SDS Running Buffer (both by Invitrogen™ NuPAGE). The gels were then electrotransferred onto PVDF membranes (Invitrolon™, Novex™, LC2005) via wet transfer. Membranes were blocked using 5% (w/v) not-fat dry milk in 0.1% (v/v) tween-20 in PBS (PBST) for at least 40 min at room temperature, and then incubated overnight at 4°C with the different primary antibodies (anti-gluA1 ser845, PhosphoSolutions, p1160-845; anti-GluA1, Santa Cruz, sc-55509; anti-GAPDH, Sant Cruz, sc-32233). Bound antibodies were detected using horseradish peroxidase conjugated Anti-rabbit and Anti-mouse secondary antibodies (mouse anti-rabbit IgG-HRP, Sant Cruz, sc-2357; m-IgGκ BP-HRP, Santa Cruz, sc-516102.), and the uniformity of sample loading was confirmed via Ponceau S staining and immunodetection of GAPDH, used as a loading control. Typhoon 9410 (Amersham Biosciences) was used to obtain digital images through chemiluminescence reaction (ECL reagents, Amersham). A semi-quantitative measurement of the band intensity was conducted using ImageJ software (https://imagej.net/ij/) and expressed as a ratio of band intensity with respect to the loading control.

#### Histology and probe localization

Following the final experimental session, all mice with bilateral silicon probe implants were anesthetized (isoflurane 2-3%) and intracardially perfused with phosphate buffer solution (PBS) and 4% paraformaldehyde (PFA) in PBS. Brains were extracted, post-fixed overnight, cryoprotected by exposure to increasing concentration of sucrose in PBS at 4°C and then sliced in coronal sections (40-50 μm thick) using a cryostat (Thermo Fisher Scientific; CryoStarTM NX50). After mounting onto slides, sections were left to dry overnight, mounted and coverslipped with standard mounting medium for fluorescence microscopy (SouthernBiotechTM; Fluoromount-G), and imaged with an upright epifluorescence microscope (Leica; DM2500). Using the acquired images, probes were identified, labelled, and registered to the Allen Mouse Brain Common Coordinate Framework^76^ using the openly available histological software ‘Histological E-data Registration in rodent Brain Spaces’ (HERBS)^77^. This allowed us to obtain a precise position for every channel in the common coordinate framework space. Slight adjustments or alignments between probes were further made using electrophysiological features such as current source densities computed on averages potentials time locked to ON-period initiations^78,79^.

#### Floor texture recognition task

A floor texture recognition (FTR) task was used to assess memory consolidation. The arena consisted of an acrylic chamber 53 cm in length by 32 cm in width, with black walls 35cm in height. A USB camera mounted above the arena was used to record the mouse’s position. During both testing and acquisition, an identical and simple cylindrical object (a small glass bottle filled with colored stones) was placed on both sides of the arena. All floors used were constituted of clear polycarbonate plastic, which sat on top of a white acrylic background. The floor was split into two halves along the long axis, and as such each half of the floor (size 32 cm x 26.5 cm) consisted of a clear polycarbonate sheet (∼ 6 mm thick) with either a smooth finish or a pebbled finish (both materials from ePlastics®). Experiments were counterbalanced so that half of subjects were exposed to two smooth floors during the acquisition and then the pebbled floor served as the novel texture in the test, and half of the subjects were exposed to two pebbled floors during the acquisition and the smooth floor served as the novel texture during the test.

All SOM+ mice used in the FTR task were habituated to handling for 7 days prior to the acquisition phase of the test. Habituation consisted of placing a gloved hand in the mouse’s home cage for 10-15 minutes per day, during which time the mouse was allowed to freely explore, as has been shown to be critical in other novelty-recognition tasks of memory consolidation^80^.

Mice were monitored by video in their home cage to ensure that sleep patterns were regular, and to ensure that mice in the sleep condition met a minimum criterion of 30% time spent asleep (mean = 45.5%, SD = 14.1%) during the 1 hour when other mice were being sleep deprived. To accurately estimate total sleep time based on video alone, we created a pose tracking model using DeepLabCut (DLC^37^), using a large number of frames across a wide variety of different cameras and mice, all from an overhead view of the mouse’s home cage. The model was trained to predict the position of a number of ‘nodes’ on the mouse visible from overhead, including the nose, head, ears, center, tail, shoulders, and hips. After training, the model was able to accurately label the position of these nodes from an overhead shot of the mouse. We used this model to predict each mouse’s position on every frame (after down sampling to 1 frame/second) of a video which started the day before acquisition and continued until after testing was completed. We then used these predictions to compute a form of actigraphy (‘DLC-actigraphy’, Extended Data Fig. 7) that could accurately estimate a mouse’s total sleep time. After getting the X-Y prediction across all frames for all nodes, we computed for each node the movement vector across all frame changes (i.e. where ΔX and ΔY represent the change in the X and Y position of a node from one frame to the next, the movement vector is equal to (ΔX^2^ + ΔY^2^)). The movement was then averaged across all nodes to get the average movement of the mouse, and that averaged movement was robust Z-scored. A threshold was set at (-mean/std) of the Z-scored movement, and any times falling below that value were labelled as sleep, while times above that value were labelled as wake. We validated the ability of the DLC-actigraphy method to predict total sleep time using overhead videos of mice that also had synchronized EEG or LFP and EMG recordings, and that had manually labelled vigilance states, which served as ground truth. Across 21 recordings from 11 individual subjects with good overhead recordings, which constituted ∼395 hours of total recording time, this method predicted total sleep time for most recordings within 5% of the manually labelled hypnogram, with an average error (compared to the manual hypnogram) of 0.55% (Extended Data Fig. 7d,e). We used a similar but separate DeepLabCut model to track the position of the mice in the arena, which produced highly accurate and reliable position predictions likely due at least in part to the constrained environment and low variance of frames to be labelled. All labelling of each mouse’s position and quantification of arena position and novelty ratios associated with the FTR task were based on the unmodified predictions of this model.

On the day of acquisition, at the lights-on time, mice were placed into the arena with two identical floor textures and left to freely explore for 15 minutes. To reduce behavioral variability during the acquisition phase, only mice that spent at least 1/3 of the total acquisition time (i.e. 5 minutes) on both sides of the arena were used for the full experiment. After the acquisition phase, these mice were removed from the arena, placed back into their home cage, and either allowed to sleep, sleep deprived, or sleep deprived concurrently with OFF period induction for one hour. For the induction, to mimic real NREM sleep as closely as possible, the temporal sequence of OFF periods used to drive the laser matched exactly (i.e. with identical timings and durations) the one previously recorded during the first hour of recovery NREM sleep following SD on a control probe of a mouse with a very high-quality implant in M2. Laser power at the ferrule tip in these experiments was similar to that used in the synaptoneurosome experiments. 24 hours following the start of the acquisition phase, mice were placed back into the arena again for 15 minutes. Using the model predictions of the mouse’s nose position, the novelty ratio during the 5-minute testing phase (first 5 minutes of exploration during the test) was quantified and expressed as the ratio of time on the side of the arena with the novel floor texture compared to the total time in the arena during the testing phase (5 minutes).

## Statistical analysis

The results of every statistical test, linked to the corresponding figure and panel, are reported in Extended Data Table 1. The table reports the type of test run, the test statistic, the *p*-value, the effect size method and value, and *N* for all statistical tests performed. All statistics were implemented in python, largely using the SciPy and Pingouin statistical libraries^81,82^. Statistical source data and code for reproduction of statistical tests and plots are available at the following repository: (https://github.com/CSC-UW/OFF_PERIOD_INDUCTION_MATERIALS). No valid subjects were selectively excluded from any analyses, except in the case of a single ACR mouse which was included in bandpower analyses due to the presence of high-quality LFPs, but due to insufficient MUA spiking activity was excluded from other electrophysiological analyses relying on OFF period detection or measuring spike-time synchrony. Excluding this mouse entirely did not affect the results of any relevant analysis.

All significance testing employed in this paper was two-tailed with α = 0.05. For all significance tests where multiple comparisons were not necessary, normality was first assessed using a Shapiro-Wilk test, and the appropriate parametric or non-parametric tests were performed. Specifically, when data did not violate parametric assumptions per the Shapiro-Wilk test (*P* > 0.05), a paired or unpaired t-test was performed as appropriate; otherwise, the corresponding non-parametric test was used (Wilcoxon signed-rank test or Mann-Whitney U test, respectively). If the same null hypothesis was evaluated across e.g. channels or frequency bins, *p*-values were adjusted with the Benjamini–Hochberg false-discovery-rate procedure.

Sample sizes were not predetermined using statistical methods, but are in line with previously published similar work^16,17,23,74^ and sample sizes used across all experiments are reported in Extended Data Table 1 and in the main text.

In the case of multiple group comparisons, normality was assessed for all components using Shapiro-Wilk, and equal variance was assessed using Levene’s test of homoscedasticity. If normality and homoscedasticity assumptions were met, one- or two-way ANOVA tests were performed as appropriate. Significant omnibus effects were followed with pairwise post-hoc contrasts using Tukey’s HSD.

## Data availability

All data and code required to reproduce all statistical tests and plots in the paper is available at the following repository: (https://github.com/CSC-UW/OFF_PERIOD_INDUCTION_MATERIALS). Any further materials will be made available upon reasonable request.

## Acknowledgments

We would like to thank all members of the Center for Sleep and Consciousness at the University of Wisconsin - Madison for helpful discussions and thoughtful input. We are especially grateful to Claire Punke for extensive help with sleep scoring, sleep deprivation and data acquisition. Additionally, we would like to thank Nate Fischer, Kade Hagen, Alex Clark, Lily Reardon, Diella Zhubi, and Johanna Ellefson for help with histology, colony management, sleep scoring, and sleep deprivation.

## Author contributions

Conceptualization: KD, GT, CC Methodology: KD, FS Visualization: KD, FS

Funding acquisition: GT, CC Writing: KD, FS, GT, CC

## Extended Data

**Extended Data Fig. 1.**
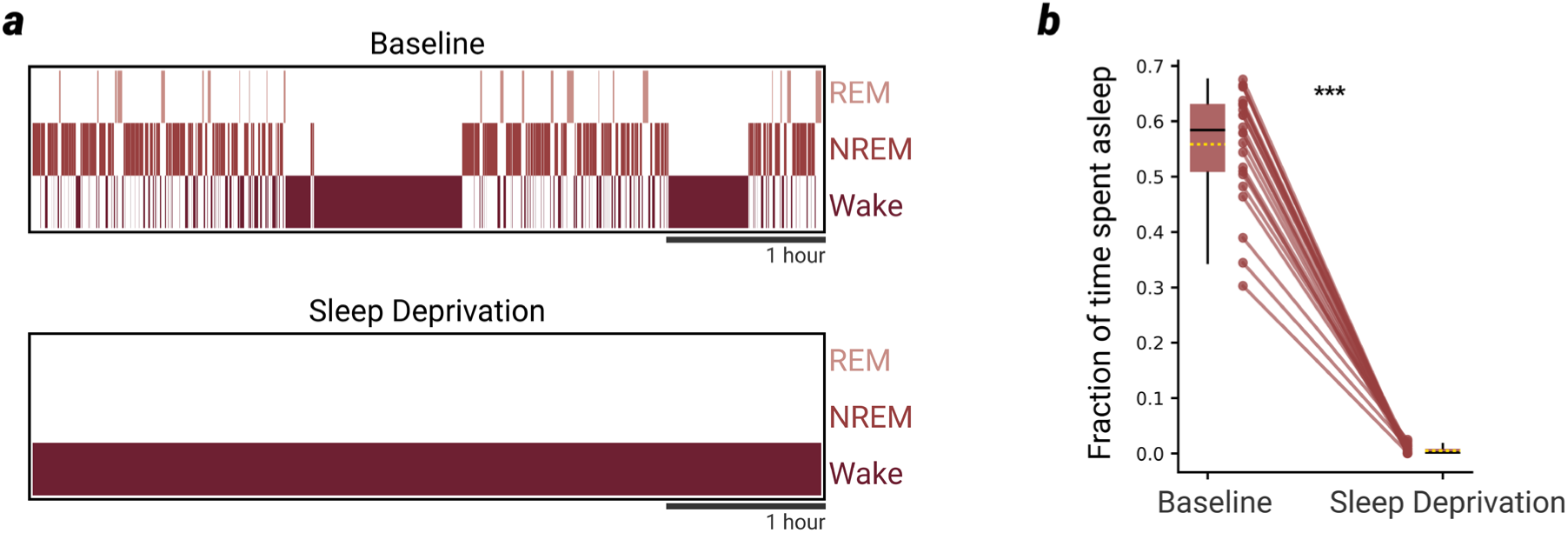
Effectiveness of sleep deprivation (SD) by exposure to novel objects **a**, Example hypnogram during the first 5 hours of the 24-hour baseline day (top), and the 5-hour SD the following day (bottom). **b**, Fraction of time spent asleep during the first 5 hours of the baseline and during the 5-hour SD across all OFF period induction experiments for SOM+, ACR, and control mice (n=24).

**Extended Data Fig. 2.**
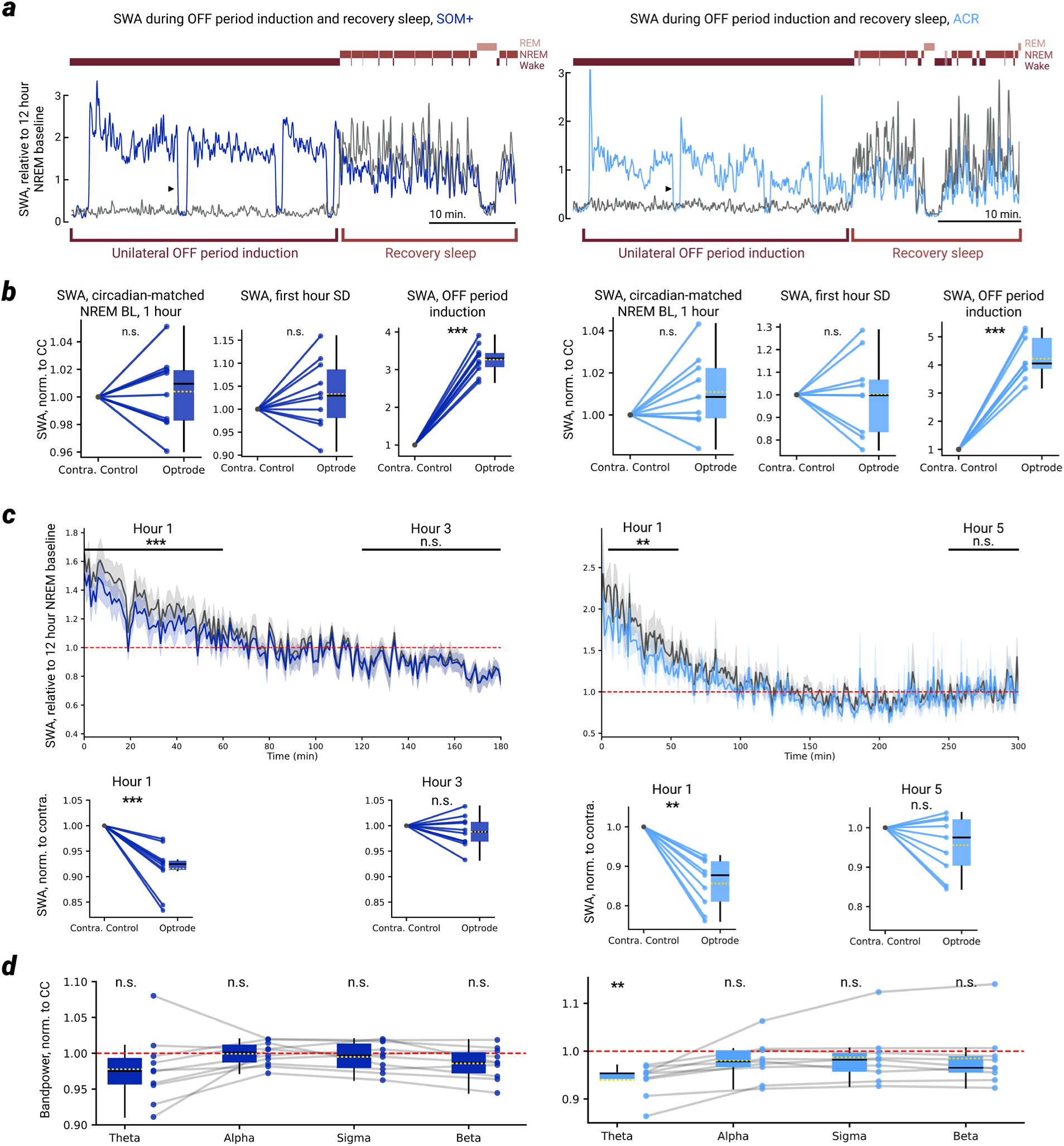
SWA dynamics following 1-hour NREM recovery **a**, SWA over the course of OFF period induction, and at the start of the NREM recovery period, for an example SOM+ and ACR mouse. Black triangle indicates the first 30-45 second break in between bouts of OFF induction (see methods). **b**, SWA during the first 1 hour of NREM sleep in the baseline (left), during the first 1 hour of SD (middle), and during the entirety of the 30-min OFF period induction period (right) at the end of SD, for all SOM+ and ACR mice. Note the tight correspondence in SWA between optrode and contralateral control during both baseline sleep and early (first hour) SD, and the extreme dissociation during OFF period induction. **c**, NREM SWA over the course of a few hours following SD across all SOM+ and ACR mice. Only NREM data are shown, relative to start of recovery sleep. On average, the optrode SWA renormalizes to the contralateral control levels within ∼3 hours for SOM+ mice, and ∼5 hours for ACR mice. Horizontal bars above line plot represent the 1 hour period of data taken for the corresponding statistical plots beneath. **d**, Bandpower on the optrode (normalized to the contralateral control level) during the 1 hour NREM recovery period, across theta (4-8 Hz), alpha (8-13 Hz), sigma (11-16 Hz), and beta (13-30 Hz), for all SOM+ and ACR mice. Red dotted line represents the contralateral control probe. In all panels, *P<.05 | **P<.005 | ***P<.0005 | n.s. P>0.05.

**Extended Data Fig. 3.**
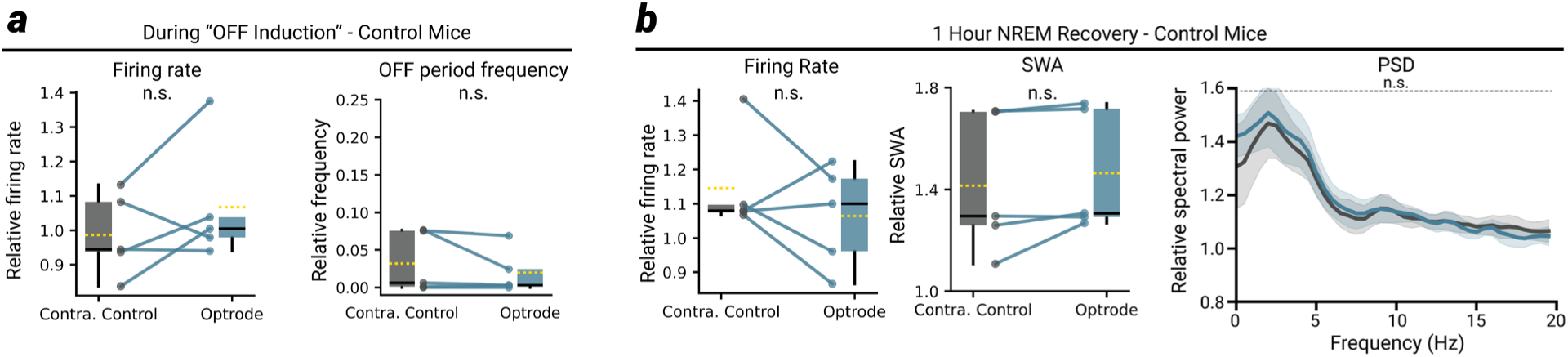
Control experiments in wildtype mice **a**, No change between optrode and contralateral control probe in either overall firing rate (left, both probes relative to first hour of SD) or OFF period frequency (right, both probes relative to 12-hour NREM baseline mean) when the OFF period induction protocol is performed on wildtype control mice. **b**, During the 1-hour NREM recovery period following SD, neither the overall MUA firing rate, nor the SWA level, nor the average power spectral density (PSD) differs between optrode and contralateral control probe. All plots express data relative to the 12-hour NREM baseline mean. In all panels, *P<.05 | **P<.005 | ***P<.0005 | n.s. P>0.05.

**Extended Data Fig. 4.**
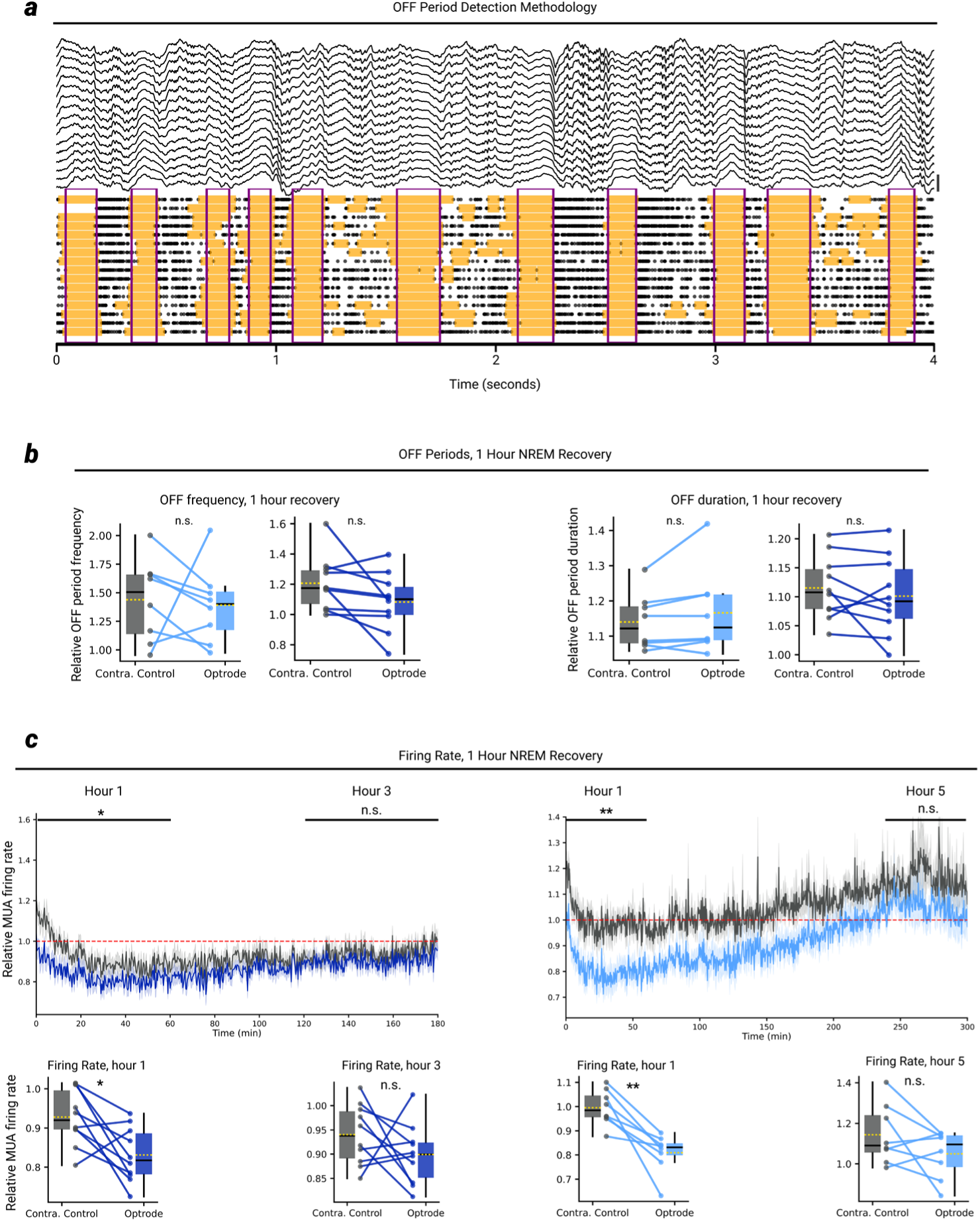
OFF periods and firing rates during the 1-hour NREM recovery period **a**, OFF period detection method. Shown is 4 seconds of example data from the NREM recovery period of a representative mouse. OFF periods are first detected at the single-channel level (orange shading) using MUA data, and only when most channels simultaneously enter an OFF period for at 50 ms is a global OFF period detected (purple boxes; see materials and methods). Scale bar at right represents 1 mV. **b**, OFF period frequency (left) and duration (right) during the 1-hour NREM recovery period across all SOM+ and ACR mice. **c**, NREM overall MUA firing rate over the course of a few hours following sleep deprivation across all SOM+ and ACR mice. Only NREM data are shown, relative to start of recovery sleep. On average, the optrode firing rate renormalizes to the contralateral control levels within ∼3 hours for SOM+ mice, and ∼4-5 hours for ACR mice. Horizontal lines above plot represent the 1-hour period of data taken for the corresponding statistical plots beneath. In all panels, *P<.05 | **P<.005 | ***P<.0005 | n.s. P>0.05.

**Extended Data Fig. 5.**
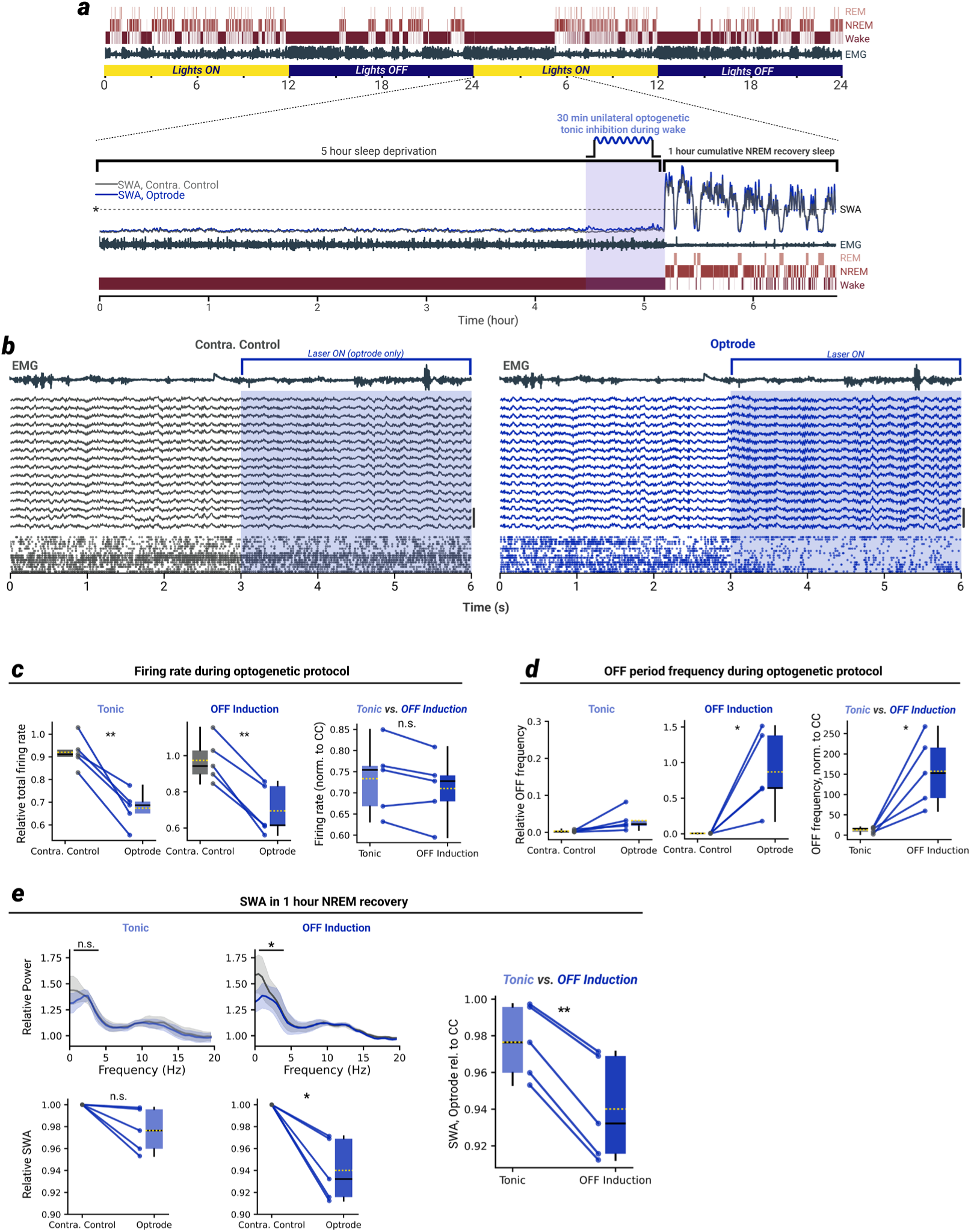
Tonic inhibition is ineffective in releasing sleep pressure **a**, Experiment overview. In a subset of SOM+ mice, unilateral tonic inhibition of overall MUA firing rate was performed for the final ∼30 minutes of a 5-hour SD. All other aspects of the experimental design were identical to the OFF period induction experiments (see materials and methods). *Dotted line indicates the 12-hour NREM baseline SWA mean. **b**, 6 seconds of raw data from a representative mouse showing tonic inhibition of MUA using continuous activation of SOM+ neurons on the optrode, and not on the contralateral control probe (see materials and methods for details). Scale bars at right of each plot represent 2 mV. **c**, Changes in overall MUA firing rate during tonic inhibition (left) and OFF period induction (middle), and direct comparison between the two optogenetic protocols (right). Left and middle panels display data relative to the mean firing rate from the first hour of SD, right panel shows optrode data only, relative to the contralateral control probe. **d**, As in **c**, for changes in the frequency of OFF periods. Left and middle panels display data relative to the 12-hour NREM baseline mean, right panel shows optrode data only, relative to the contralateral control probe. **e**, PSD plots (top) and SWA in the 1-hour NREM rebound following tonic inhibition (left) and OFF period induction (middle). Large plot at right shows that, for every animal tested, OFF period induction was more effective in releasing sleep pressure following SD than tonic inhibition (plot shows only optrode data, relative to contralateral control probe). In all panels, *P<.05 | **P<.005 | ***P<.0005 | n.s. P>0.05.

**Extended Data Fig. 6.**
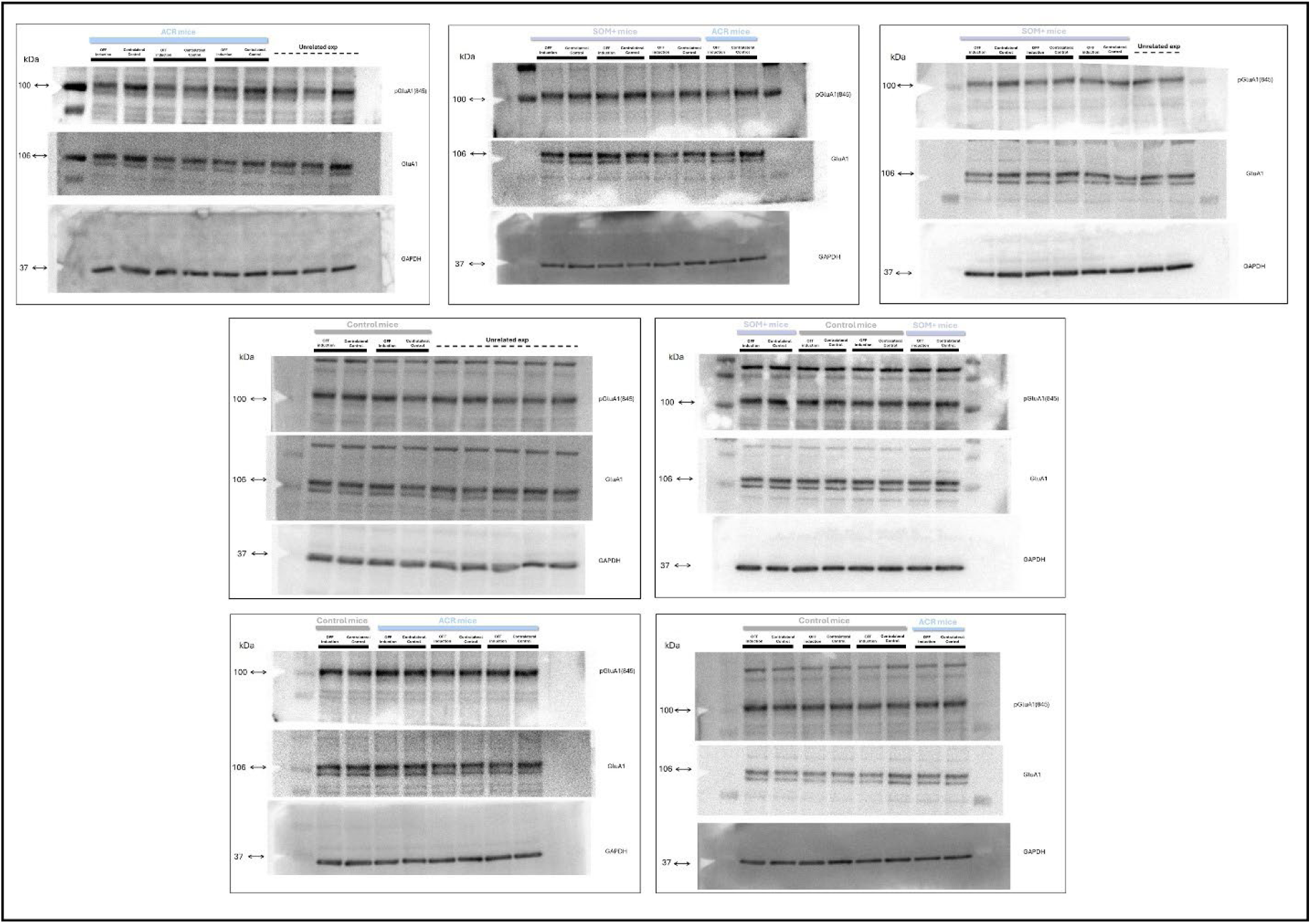
Uncropped Western blot membranes Uncropped, full-length images of Western blot membranes. Membranes were cut around 50 kDa to allow probing with multiple antibodies. For each membrane, the top image shows the pGluA1(845) bands, the middle image shows the GluA1 bands, and the bottom image shows the GAPDH housekeeping bands. Molecular weights are indicated on the left side of each image: 100 kDa for pGluA1(845), 106 kDa for GluA1, and 37 kDa for GAPDH. For each subject, the OFF-induction side sample was loaded first, followed by the corresponding contralateral control side sample, as indicated by the black bars above each membrane lane. For experimental condition information: the light blue bars indicate ACR mice, periwinkle bars indicate SOM+ mice, gray bars indicate Control mice, and dotted lines denote unrelated experimental data.

**Extended Data Fig. 7.**
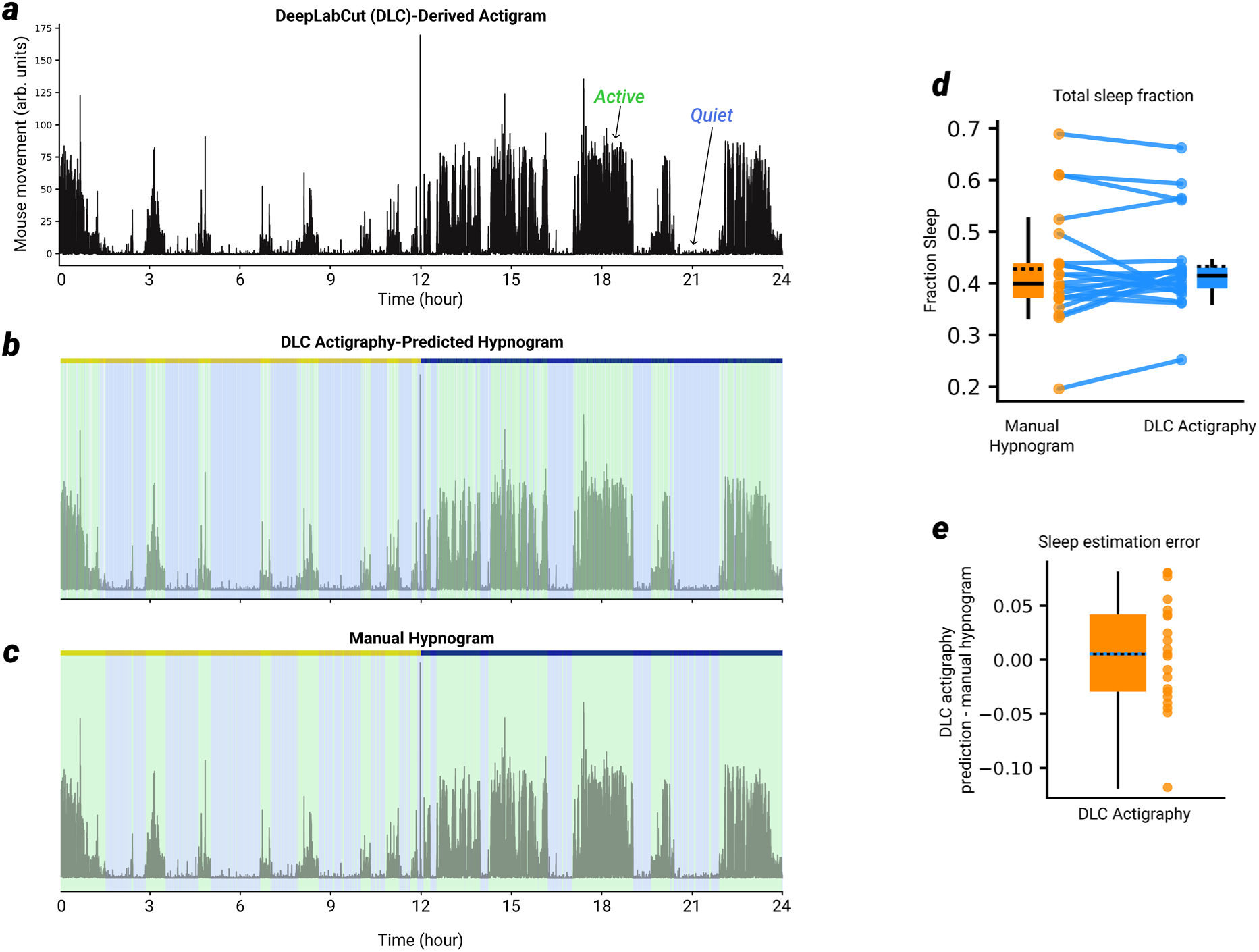
Validation of actigraphy for accurate estimation of total sleep time **a**, Actigram for a single example mouse, over a 24-hour video recording. In addition to video recording, this example mouse (as for all mice in this figure) was implanted with EMGs and at least one silicon probe. Therefore, the DeepLabCut-based (DLC) actigraphy method of total sleep estimation could be compared to ground-truth manual sleep scoring (see materials and methods). **b**, Hypnogram predicted by the DLC actigraphy method described in the materials and methods (which requires only overhead video) Green shading indicates wake, blue shading indicates sleep. **c**, Manual hypnogram, scored on the basis of LFP and EMG signals. **d**, For each recording in the testing set, shown is the total fraction of time spent asleep as judged by manual scoring based on EMG and LFP signals (orange) or by DLC actigraphy (blue). **e**, DLC-actigraphy error for each recording in the testing set, defined as the total sleep fraction prediction made by DLC actigraphy – the total sleep fraction according to the manually-scored hypnogram.

---

**Extended Data Table 1.**
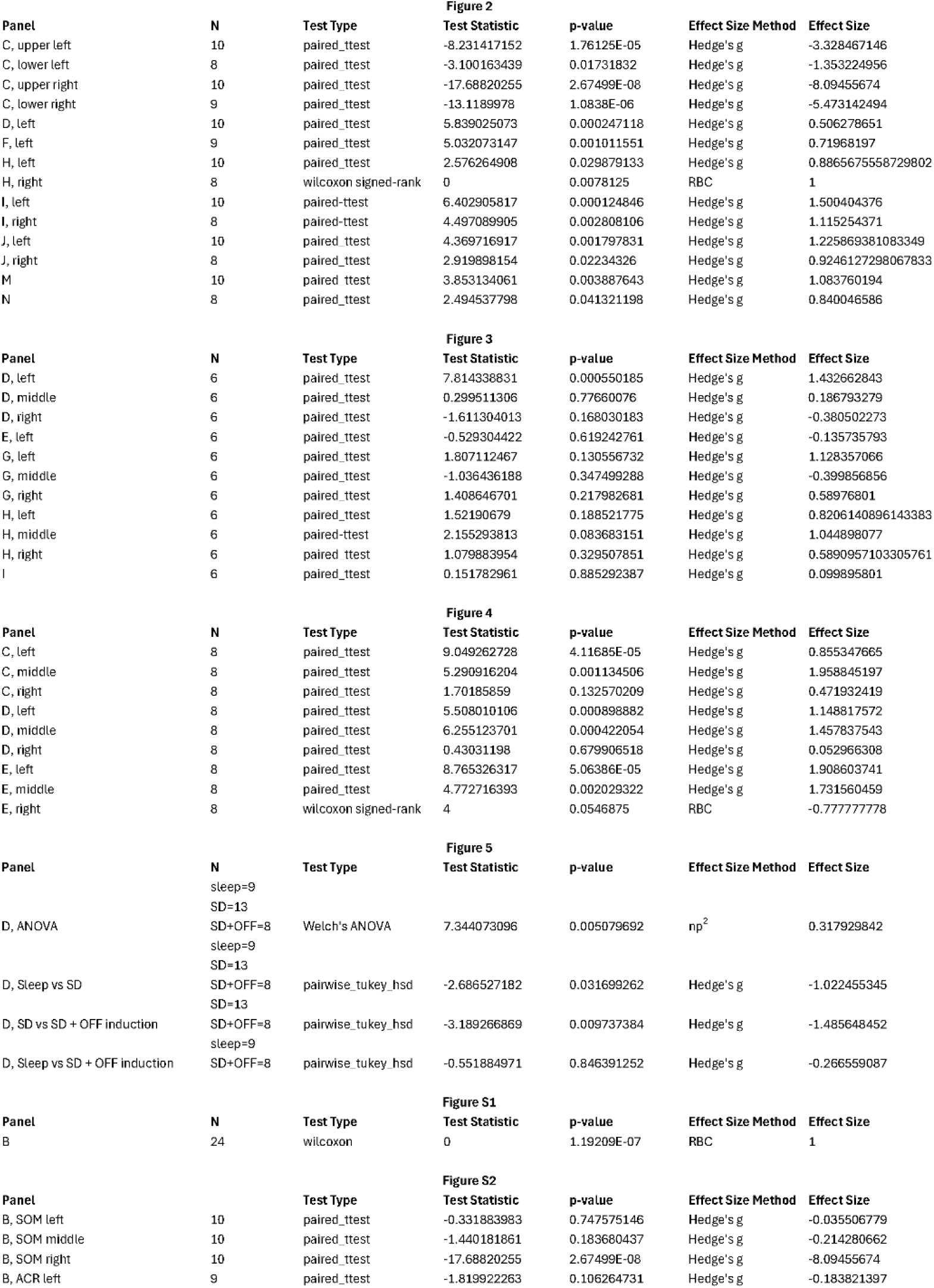
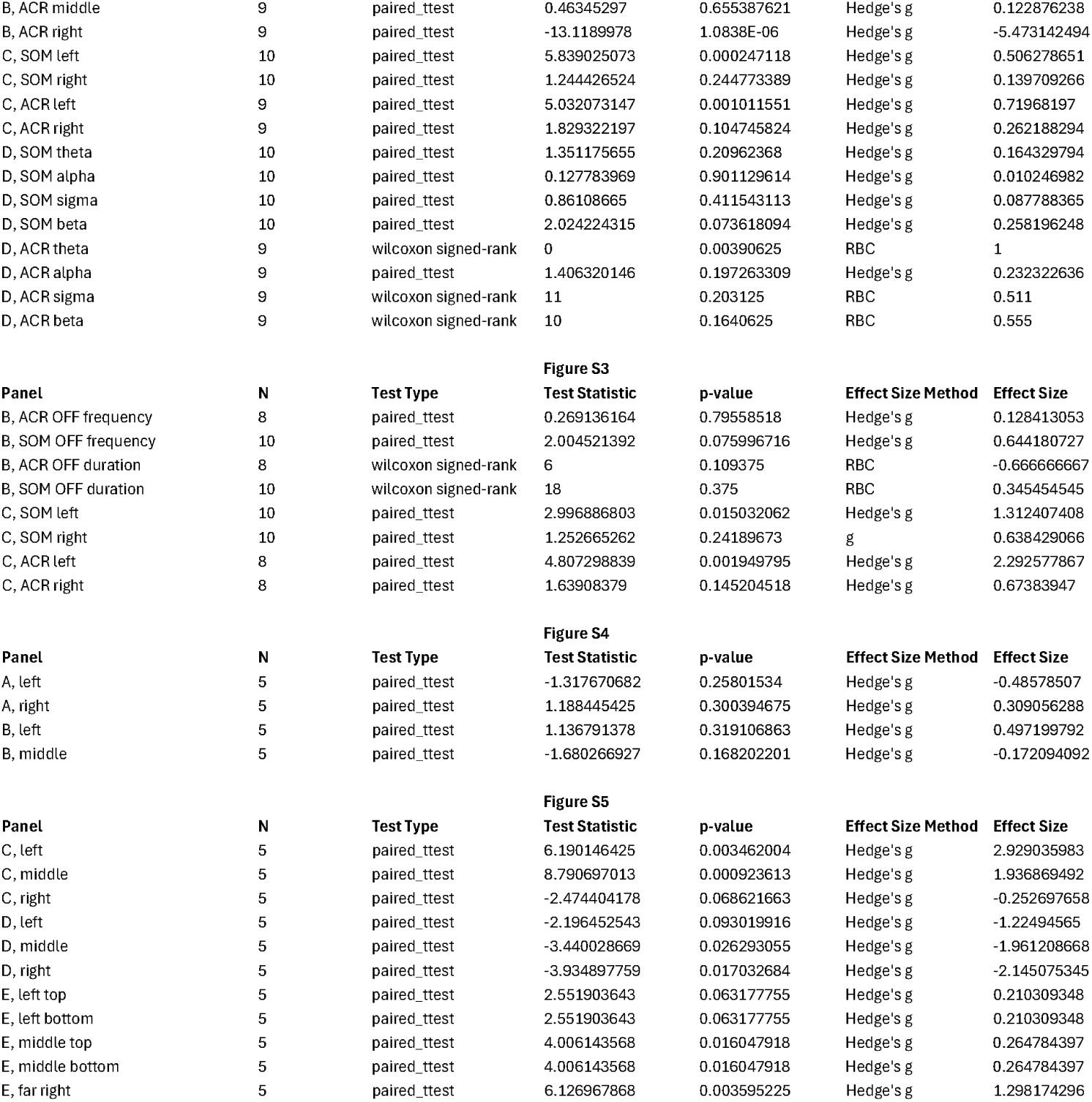
Statistical summary Complete statistical information for all statistical tests run in this paper, with corresponding figure and panel labels. Reported for each test is number of mice, the test type, the test statistic, the *p*-value, and the effect size method and value. Note that complete statistical tables across all frequencies for power spectral densities in figures 2, 3, and S3 can be found with the accompanying statistical source data.

**Extended Data Movie 1** Demonstration of OFF period induction during wake Movie shows 10 seconds of raw data (LFP and MUA) as animal is awake during SD (left), in NREM sleep (middle), and is awake with concurrent OFF period induction (right). Note the dissociation between the two probes during OFF period induction, and that during OFF period induction the animal remains awake and interacts with the environment.

